# A Moonlighting Function of Kinetochore Complex Proteins in Golgi Assembly of Neural Stem Cells

**DOI:** 10.64898/2026.05.24.727544

**Authors:** Mahekta R. Gujar, Htet Yamin Aung, Yang Gao, Dongliang Ma, Divya Suresh, Jiaen Lin, Xiang Teng, Yusuke Toyama, Fengwei Yu, Hongyan Wang

**Author notes:** Lead contact: Hongyan Wang.

## Abstract

Kinetochores play an essential role in mitosis, and human kinetochore variants are associated with microcephaly. However, their functions in quiescent/interphase cells are unknown. Here, we demonstrate that *Drosophila* kinetochore proteins promote the reactivation of quiescent NSCs and their regeneration upon injury by organizing Golgi-dependent acentrosomal microtubules. *Drosophila* kinetochore proteins localize to the Golgi vicinity in quiescent NSCs and are required for proper localization of Golgi proteins, including the Golgi-resident small GTPase Arf1 known to regulate quiescent NSC reactivation and regeneration. Moreover, *Drosophila* kinetochore proteins physically associate with Arf1 and function upstream of it during reactivation. Remarkably, human kinetochore protein SPC25 localizes to Golgi in interphase NSCs and HEK293T cells. Furthermore, human kinetochores have a conserved function in promoting Golgi organization and, in turn, microtubule dynamics, in interphase cells. Together, our findings reveal a conserved moonlighting role for kinetochores in Golgi assembly in quiescent *Drosophila* NSCs and human interphase cells.

## Introduction

Neural stem cells (NSCs) are vital for nervous system development, regeneration, and repair. In the adult mammalian brain, most NSCs exist in a quiescent or mitotically dormant state characterized by a lack of proliferation or differentiation allowing for the limited pool of NSCs to be preserved ^1, 2^. These quiescent NSCs (qNSCs) can reenter the cell cycle and resume their proliferation, in a process named reactivation to generate new neurons in response to physiological stimuli such as injury, availability of nutrients, or physical exercise ^3^. Conversely, stress, ageing, or anxiety severely reduce NSC proliferation. The capacity of NSCs to dynamically transition between quiescent and proliferative states is fundamental to adult neurogenesis and regenerative potential, and disruption of this quiescence–reactivation balance is associated with impaired neurogenesis and neurodevelopmental disorders, including microcephaly ^4, 5^. Therefore, elucidating the molecular, cellular, and environmental mechanisms that orchestrate NSC reactivation is critical for advancing our understanding of brain development and the etiologies of neurodevelopmental disorders.

The *Drosophila* larval brain provides a powerful *in vivo* model for elucidating mechanisms of NSC quiescence and reactivation. At the end of embryogenesis, *Drosophila* NSCs, also known as neuroblasts, shrink in size and enter a quiescent state, which is regulated by the combined function of temporal identity factors, homeobox genes, and the transcription factor Prospero ^6–9^. Approximately 24 hours after larval hatching (ALH), feeding and dietary amino acids trigger an evolutionary conserved insulin/IGF-like signalling cascade. The blood-brain barrier (BBB) glial cells secrete insulin-like peptides (Dilp2, Dilp6), which activate the evolutionary conserved insulin/IGF receptor (InR/IGF-1R) pathway to drive NSC reactivation ^10–12^. Similar to *Drosophila* NSCs, mammalian NSCs are also activated by IGF-1 signaling, and mutations in the human IGF-1 receptor are linked to microcephaly ^13,14^. Additionally, several intrinsic factors that regulate NSC reactivation have also been identified, such as, Hsp83/Hsp90 chaperones that assist in InR activation, spindle-matrix proteins, striatin-interacting phosphatase and kinase family proteins, E3 ubiquitin ligase CRL4^Mahj^, Pr-set7, SUMO pathway proteins, microtubule interacting proteins Msps/XMAP215, Patronin/CAMSAP and Arf1 ^7, 15–23^. In contrast, the evolutionarily conserved Hippo pathway operates to maintain NSC quiescence: activation of Hippo (Hpo/Warts) suppresses growth and proliferation ^24–26^. Thus, NSC reactivation requires coordination between extrinsic signals (nutrition, glial-secreted insulin-like peptides) and intrinsic regulatory machinery.

A distinct morphological hallmark of quiescent *Drosophila* NSCs is the presence of a primary cellular protrusion projecting toward the neuropil ^12, 27^. This protrusion is enriched in acentrosomal microtubules oriented predominantly plus-end out, similar to neuronal axons, while the centrosomes in newly hatched larvae are immature and lack robust microtubule nucleation capacity ^17, 21^. Our recent work has revealed that the Golgi apparatus acts as an acentrosomal microtubule organizing center (MTOC) in these qNSCs ^19^. Remarkably, quiescent NSC cellular protrusions can be regenerated upon injury by laser severing, and this regeneration relies on Golgi and microtubule growth in qNSCs ^19^. Two essential Golgi-associated proteins, the small GTPase Arf1 and its guanine nucleotide exchange factor Sec71/ArfGEF are required for both NSC reactivation and regeneration, by regulating microtubule growth ^19^. Msps/XMAP215 a key microtubule polymerase functions downstream of Arf1 in promoting microtubule polymerization, whereas the microtubule minus-end binding protein Patronin/CAMSAP acts as a pivotal upstream regulator of this process ^19, 20^. Collectively, Patronin-Arf1-Msps pathway is necessary for both reactivation of NSCs and for regeneration of cellular protrusions after injury ^19, 20^.

The kinetochore is a specialized structure formed during mitosis that connects chromosomes to spindle microtubules, ensuring correct segregation of genetic material into daughter cells, a function that has been widely conserved across eukaryotes. The Kinetochore complex resides at the centromere of each chromosome and, by capturing spindle microtubules and attaching them to chromosomes, allow the spindle to pull chromosomes toward the poles ^28–30^. One subcomplex, the inner kinetochore, is bound to centromeric heterochromatin while three other subcomplexes, composed of the Knl1 (also called Spc105 in *Drosophila*), Mis12, and Ndc80 complexes, known as the KMN network, bridges inner centromere elements to the outer microtubule-binding machinery ^28, 29, 31, 32^. In *Drosophila*, inner centromere proteins (Cenp-A, Cenp-C) and Mis12 localize to centromeres continuously, both in interphase and mitosis ^33^. By contrast, other kinetochore components, especially those of the Knl1 and Ndc80 complexes, are recruited only during mitosis ^33^. Once fully assembled, outer kinetochore proteins such as Nuf2 and Ndc80 capture the ends of growing microtubules during mitosis, enabling chromosome segregation ^30, 34–36^. Structural studies have revealed that the elongated Ndc80 complex directly binds spindle microtubules through calponin-homology domains and flexible N-terminal tails, allowing kinetochores to establish robust attachments to dynamic microtubule plus ends ^37–40^. These attachments are tightly regulated by Aurora B kinase-mediated phosphorylation, which destabilizes improper kinetochore–microtubule interactions and ensures faithful chromosome segregation ^41–43^. Although the kinetochore and its regulatory complexes are classically associated with dividing cells, most of them have not been strongly tied to functions outside cell division. Recent studies, however, point to roles in postmitotic (non-dividing) cells, particularly neurons ^44–46^. For example, in *C. elegans*, components of the KMN network localize in ciliated dendrites of amphid neurons and contribute to their extension^44^. In *Drosophila*, mutants of Mis12 affect neuromuscular junction (synaptic) structure, and knocking down other KMN proteins or the centromeric histone Cid/CENP-A has similar effects ^45^. In rodent hippocampal neurons, reducing Mis12 increased dendritic protrusions ^45^, suggesting that kinetochore proteins appear to influence neuronal form across species. Furthermore, mutations in kinetochore proteins have been shown to cause brain disorders, particularly neurodevelopmental disorders such as microcephaly ^47–49^. Therefore, understanding whether and how the kinetochore complex proteins regulate NSC reactivation will provide insights into developing therapeutic targets for neurodevelopmental disorders.

In this study, we investigated the non-canonical function of kinetochore proteins in quiescent/interphase cells. We show that kinetochore proteins are important for the reactivation and regeneration of qNSCs, as well as proper localization and function of key Golgi proteins Arf1 and Sec71/Arf1GEF, to regulate acentrosomal microtubule growth in the cellular protrusion of qNSCs. Moreover, kinetochore proteins can physically associate with Arf1. Genetic analyses support our model that the kinetochore complex functions upstream of Arf1 and its effector Msps/XMAP215 to regulate acentrosomal microtubule growth, NSC reactivation, and regeneration upon injury. Finally, we show that kinetochores have a conserved, moonlighting function in promoting Golgi assembly in human interphase cells.

## Results

### Kinetochore complex proteins are required for qNSC reactivation in *Drosophila*

To understand if the kinetochore complex proteins are important for qNSC reactivation, we performed EdU (5-ethynyl-2’-deoxyuridine) incorporation assay at 24 h after larval hatching (ALH). As loss of function mutants of most kinetochore proteins were embryonically lethal, we examined two presumptive loss-of-function *nuf2* alleles, *nuf2^EX^*^50^ and *nuf2^SH2040^,* that survived till 24h ALH. *nuf2^EX50^* was generated by the imprecise excision of P(lacW)Nuf2^SH2276^ which results in an 644bp deletion that removes the first half of the gene, while *nuf2^SH2040^* was generated by insertion of a transposable element. 33.2% of NSCs in *nuf2^EX50^* and 31.6% of NSCs in *nuf2 ^SH2040^* failed to incorporate EdU, compared with only 10.5% of NSCs without EdU incorporation in wild-type (Figure 1A-B), Moreover, the EdU incorporation defects in *nuf2^EX50^*were fully restored by expressing a genomic Nuf2 (gNuf2) (Figure 1A-B). These observations indicate that Nuf2 is required for qNSC reactivation in the *Drosophila* larval brain.

**Figure 1.**
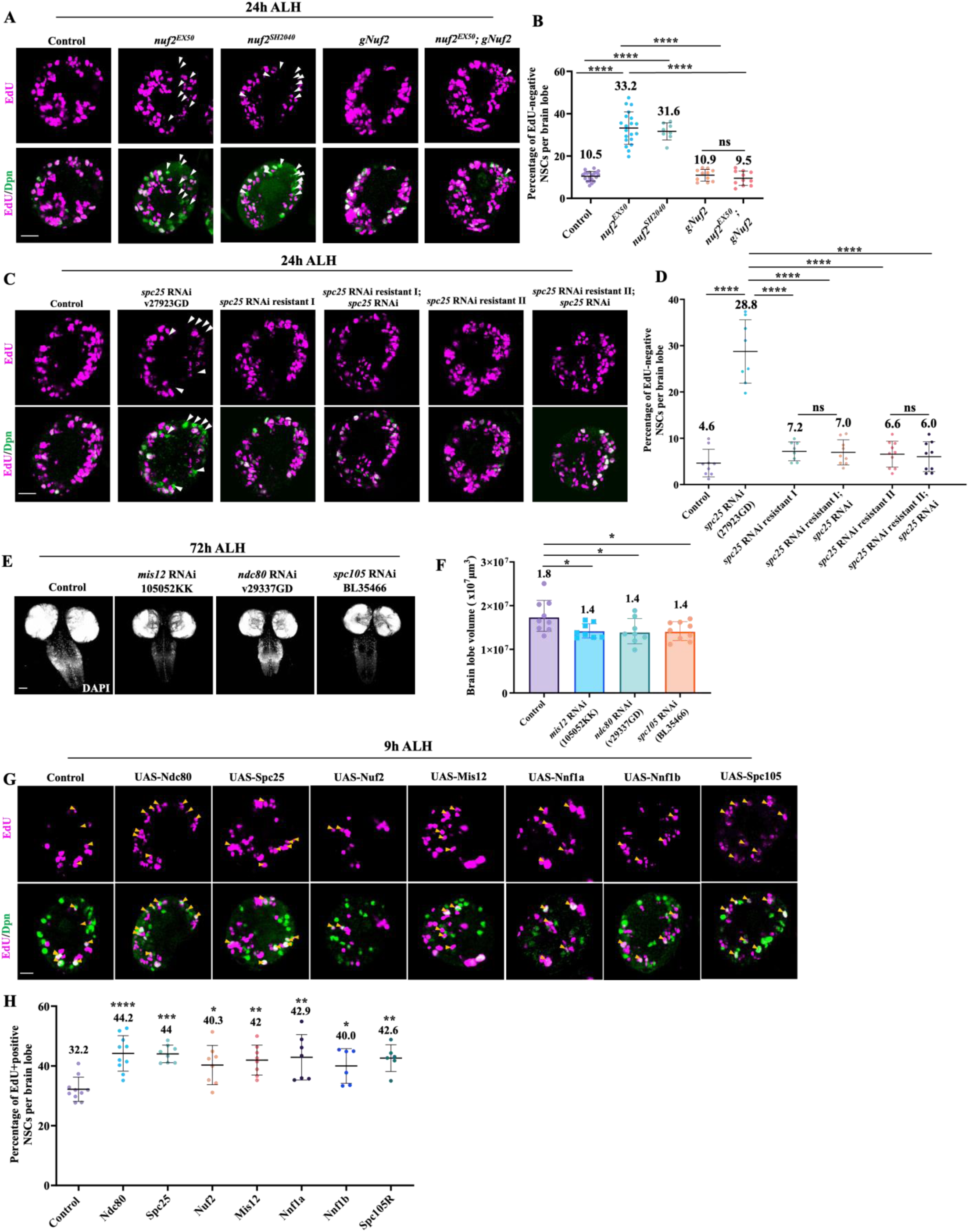
The kinetochore complex proteins are required for quiescent NSC reactivation. A) Larval brains at 24h ALH from control, *nuf2^EX50^*, *nuf2^SH2040^*, *gNuf2* and *nuf2^EX50^; gNuf2* were analysed for EdU incorporation. NSCs were marked by Dpn. B) Quantification graph of EdU-negative NSCs per brain lobe for genotypes in (A). Control, 10.5%, n=20 BL; n*uf2^EX50^*, 33.2%, n=20 BL; *nuf2^SH2040^*, 31.6%, n=8 BL; *gNuf2*, 10.9%, n=11 BL and *nuf2^EX50^; gNuf2*, 9.5%, n=11 BL. C) Larval brains at 24h ALH from control (*grh*-Gal4; UAS*-dicer2/*UAS*-β-Gal* RNAi), *spc25* RNAi (VDRC 27923GD), *spc25* RNAi (104213KK) resistant I, *spc25* RNAi (27923GD); *spc25* RNAi resistant I, *spc25* RNAi (104213KK) resistant II and *spc25* RNAi (27923GD); *spc25* RNAi resistant II, were analysed for EdU incorporation. NSCs were marked by Dpn. D) Quantification graph of EdU-negative NSCs per brain lobe for genotypes in (C). Control, 4.6%, n=9 BL; *spc25* RNAi (27923GD), 28.8%, n=7 BL; *spc25* RNAi (104213KK) resistant I, 7.2%, n=8 BL; *spc25* RNAi (27923GD); *spc25* RNAi resistant I, 7%, n=9 BL; *spc25* RNAi (104213KK) resistant II, 6.6%, n=10 BL and *spc25* RNAi (27923GD); *spc25* RNAi resistant II, 6%, n=9 BL. E) Maximum intensity z-projection of larval brains from control (*grh*-Gal4; UAS*-dicer2/*UAS*- β-Gal* RNAi), *mis12* RNAi (105052KK), *ndc80* RNAi (29337GD) and *spc105R* RNAi (BL35466), were stained with DAPI. F) Quantification of brain volume in (E). Control, 1.8 ± 0.3 x 10^7^ µm^3^, n = 9; *mis12* RNAi (105052KK), 1.4 ± 0.17 x 10^7^ µm^3^, n = 8; *ndc80* RNAi (29337GD), 1.4 ± 0.3 x 10^7^ µm^3^, n = 8 and *spc105R* RNAi (BL35466), 1.4 ± 0.2 x 10^7^ µm^3^, n = 9. G) Larval brains at 9h ALH from control (*grh*-Gal4*/*UAS*-β-Gal* RNAi), UAS*-Ndc80,* UAS*-Spc25,* UAS*-Nuf2,* UAS*-Mis12,* UAS*-Nnf1a,* UAS*-NNf1b and* UAS*-Spc105R* controlled under *grh*-Gal4 were analyzed for EdU incorporation. NSCs were marked by Dpn. H) Quantification graph of EdU+positive NSCs per brain lobe for genotypes in (G). Control, 32.2%, n=10 BL; UAS*-Ndc80*, 44.2%, n=10 BL; UAS*-Spc25,* 44%, n=8 BL; UAS*-Nuf2,* 40.3%, n=8 BL; UAS*-Mis12,* 42%, n=8 BL; UAS*-Nnf1a,* 42.9%, n=7 BL; UAS*-Nnf1b,* 40%, n=6 BL; and UAS*-Spc105R,* 42.6%, n=6 BL. Data information: (A-D) EdU incorporation was analyzed at 24h ALH by feeding larvae at 20h ALH with food supplemented with 0.2 mM EdU for 4h. White arrowheads point to NSCs without EdU incorporation. (G-H) EdU incorporation was analyzed at 9h ALH by feeding larvae at 5h ALH with food supplemented with 0.2 mM EdU for 4h. Yellow arrowheads point to NSCs with EdU incorporation. Data are presented as mean ± SD. In (B, D and H), statistical significance was determined by one-way ANOVA with multiple comparisons. *p<0.05, **p<0.01, ***p<0.001 and ****p<0.0001. Scale bars: 10μm.

Next, we examined a total of 14 RNA interference (RNAi) lines targeting various components of kinetochore proteins, including the Mis12 complex (Mis12, Kmn1, and Nnf1a), Ndc80 complex (Ndc80, Spc25, Nuf2), and Spc105R. At 24h ALH, the vast majority of control NSCs driven by the NSC specific driver *grainy head* (*grh*)*-*Gal4 were reactivated and incorporated with EdU, and only 6% of NSCs were quiescent and negative for EdU (Figure S1A-B). Remarkably, at 24h ALH, all RNAi lines, namely *mis12*, *kmn1*, *nnf1a*, *ndc80*, *spc25*, *nuf2*, and *spc105R* RNAi had significantly increased number of qNSCs indicated by lack of EdU incorporation, compared with the control (Figure S1A-B). This result suggests that various components of kinetochore complex are required for qNSC reactivation. Similarly, using Miranda (Mira) as a marker for the cellular extensions and Deadpan (Dpn) as a NSC nuclear marker, there was a significant increase in the percentage of Miranda (Mira)-positive NSCs that still extended their cellular processes (Fig S1C-D), supporting defects in qNSC reactivation in various RNAi knockdowns. To validate the efficiency of these RNAi lines in NSCs, we assessed the effects of knockdown in mitotic NSCs at 24h ALH, since the core *Drosophila* kinetochore components are well known for their function in chromosome congression. As expected, we observed strong defects in chromosome alignment and spindle abnormalities, suggesting efficient knockdowns (Figure S2A-B).

To further validate the specificity of RNAi knockdown phenotypes, we generated a synthetic RNAi-resistant transgene that encodes the wild-type Spc25 protein but bears minimal homology to the RNAi target sequence in spc25 RNAi 27923GD. The reactivation defects by EdU incorporation observed in *spc25* RNAi 27923GD larval brains at 24h ALH were fully rescued by co-expression of either of two independent RNAi-resistant *spc25* transgene lines, confirming the specificity of the *spc25* RNAi effect (Figure 1C-D).

Variants of *CASC5/Blinkin/KNL1/hSPC105*, the ortholog of *Drosophila spc105R* in humans, have been identified in patients with microcephaly (Genin 2012; Javed 2018). Remarkably, the volume of *mis12* RNAi (105052KK), *ndc80* RNAi (29337GD), and *spc105R* RNAi (BL35466) knockdown brain lobes at 72h ALH were all dramatically reduced to 1.4 × 10^7^µm^3^, compared to control larval brain volume of 1.8 × 10^7^ µm^3^ (Figure 1E-F), mimicking microcephaly phenotypes. These data indicate that kinetochore proteins are required for *Drosophila* larval brain development.

### Overexpression of kinetochore proteins triggers premature NSC reactivation

To explore the potential overexpression phenotype, we overexpressed kinetochore proteins using the NSC-specific driver *grh*-Gal4. At 9h ALH, most of the wild-type NSCs were still in a quiescent state, with about 32.2% of NSCs incorporating EdU (Figure 1G-H). Interestingly, *Ndc80*, *Spc25*, *Nuf2*, *Mis12*, *Nnf1a*, *Nnf1b* and *Spc105R* overexpression in NSCs by *grh*-Gal4 driver enhanced reactivation, evident with an increase in the number of EdU-positive NSCs (UAS*-Ndc80*, 44.2%; UAS*-Spc25*, 44%; UAS*-Nuf2*, 40.3%; UAS*-Mis12*, 42%; UAS*-Nnf1a,* 42.9%; UAS*-Nnf1b*, 40%; UAS*-Spc105R*, 42.6%), compared with 32.2% in the control (Figure 1G-H). Thus, kinetochore protein overexpression in NSCs can trigger premature NSC reactivation.

### Kinetochore proteins promote acentrosomal microtubule growth in *Drosophila* qNSCs

Kinetochore proteins have a post-mitotic function in suppressing neurite overgrowth in *Drosophila* neurons ^46, 50^, we sought to understand whether kinetochore proteins have any role in acentrosomal microtubule growth in qNSCs. At 6h ALH, the total number of EB1 comets was significantly reduced in qNSCs with a fold-change of 0.63, 0.68 and 0.72 in *mis12* RNAi 105052KK, BL35471 and BL38535, respectively (Figure 2A-B). This observation is in contrast with prior reports in neurons ^46^, suggesting that kinetochore proteins have a distinct function in promoting microtubule growth in *Drosophila* qNSCs, compared with neurons. Similarly, upon knocking down of *kmn1* (19529GD and 106889KK), *nnf1a* (18208GD and 109867KK), *ndc80* (BL33620 and 29337GD), *spc25* (104213KK and 27923GD), *nuf2* (BL35599) and *spc105R* (BL35466 and BL36100), the normalized numbers of EB1 comets were significantly reduced, as compared to 1 in the control (Figure 2A-C, Movie S1). In addition, average velocity velocities of EB1-GFP comets in most kinetochore-depleted qNSCs were significantly reduced as compared to control (Figure 2A, D-E, Movie S1).

**Figure 2.**
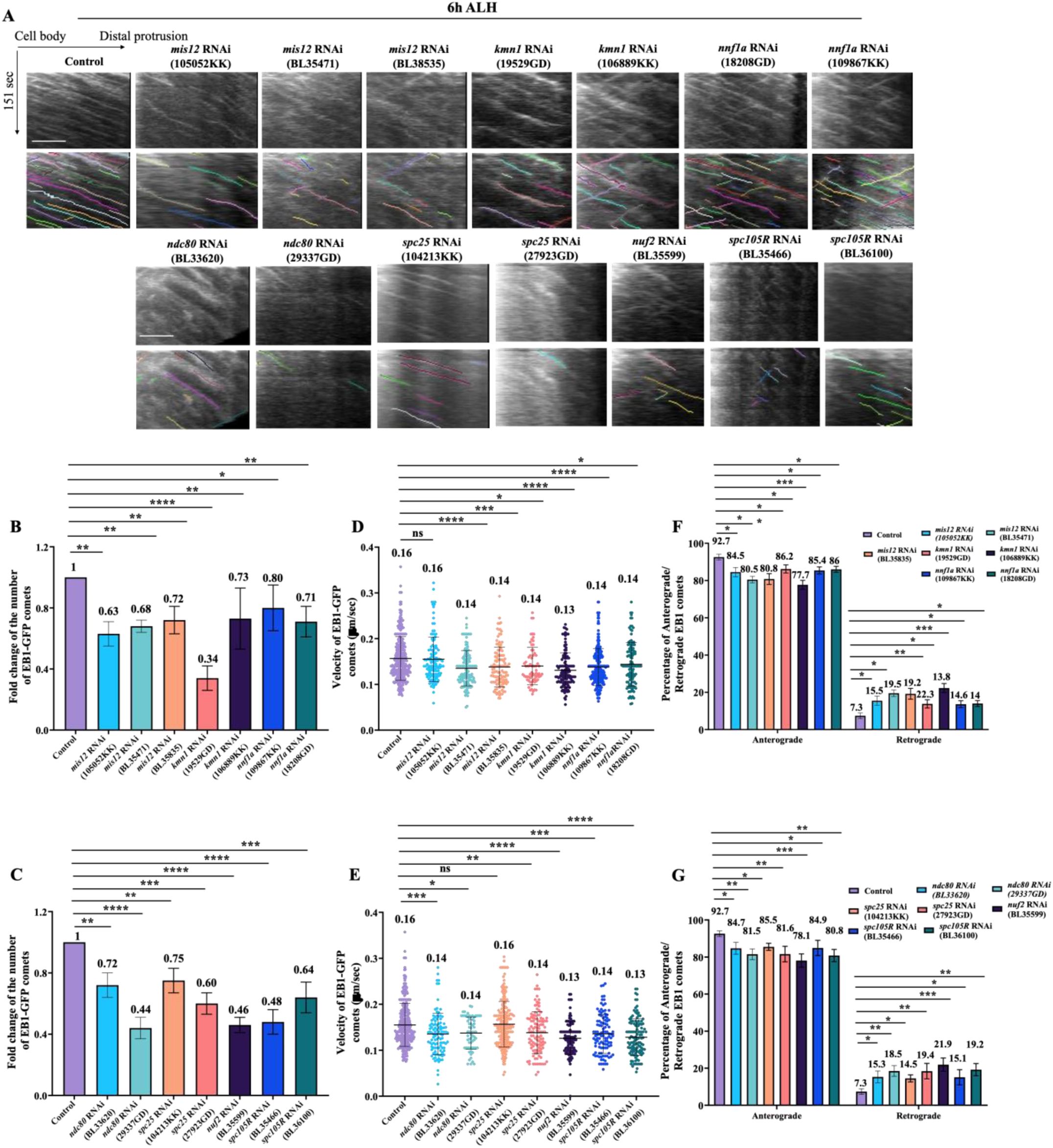
Kinetochore proteins are required for acentrosomal microtubule assembly in qNSCs. A) Kymographs of EB1-GFP comets movement in the primary protrusion of qNSCs expressing EB1-GFP under *grh*-Gal4 from the control, *mis12* RNAi (VDRC 105052KK), *mis12* RNAi (BDSC BL35471), *mis12* RNAi (BDSC BL38535), *kmn1* RNAi (VDRC 19529GD), *kmn1* RNAi (VDRC 106889KK), *nnf1a* RNAi (VDRC 18208GD), *nnf1a* RNAi (VDRC 109867KK), *ndc80* RNAi (BDSC BL33620), *ndc80* RNAi (VDRC 29337GD), *spc25* RNAi (VDRC 27923GD), *spc25* RNAi (VDRC 104213KK), *nuf2* RNAi (BDSC BL35599), *spc105R* RNAi (BDSC BL35466) and *spc105R* RNAi (BDSC BL36100) 6h ALH. The horizontal arrow indicates anterograde movement direction from cell body to the tip of the primary protrusion in qNSCs. B) Quantification graph of fold changes of number of EB1-GFP comets in the primary protrusion of qNSCs 6h ALH from various genotypes in (A). Control, 1, n=24 qNSCs, n=339 comets; *mis12* RNAi (105052KK), 0.63, n=9 qNSCs, n=128 comets; *mis12* RNAi (BL35471), 0.68, n=13 qNSCs, n=133 comets; *mis12* RNAi (BL38535), 0.72, n=11 qNSCs, n=130 comets; *kmn1* RNAi (19529GD), 0.34, n=13 qNSCs, n=68 comets; *kmn1* RNAi (106889KK), 0.73, n=11 qNSCs, n=121 comets; *nnf1a* RNAi (18208GD), 0.78, n=10 qNSCs, n=121 comets; *nnf1a* RNAi (109867KK), 0.80, n=16 qNSCs, n=219 comets. C) Quantification graph of fold changes of number of EB1-GFP comets in the primary protrusion of qNSCs 6h ALH from various genotypes in (A). Control, 1, n=24 qNSCs, n=339 comets; *ndc80* RNAi (BL33620), 0.72, n=9 qNSCs, n=106 comets; *ndc80* RNAi (29337GD), 0.44 n=15 qNSCs, n=65 comets; *spc25* RNAi (104213KK), 0.75, n=17 qNSCs, n=220 comets; *spc25* RNAi (27923GD), 0.60, n=15 qNSCs, n=110 comets; *nuf2* RNAi (BL35599), 0.46, n=12 qNSCs, n=106 comets; *spc105R* RNAi (BL35466), 0.48, n=13 qNSCs, n=106 comets and *spc105R* RNAi (BL36100) 0.64, n=11 qNSCs, n=120 comets. D) Quantification graph of velocity of EB1-GFP comets in the primary protrusion of qNSCs at 6h ALH from various genotypes in (A). Control, 0.16μm/sec, n=339 comets; *mis12* RNAi (105052KK), 0.16μm/sec, n=128 comets; *mis12* RNAi (BL35471), 0.14μm/sec, n=133 comets; *mis12* RNAi (BL38535), 0.14μm/sec, n=130 comets; *kmn1* RNAi (19529GD), 0.14μm/sec, n=65 comets; *kmn1* RNAi (106889KK), 0.13μm/sec, n=121 comets; *nnf1a* RNAi (18208GD), 0.14μm/sec, n=121 comets; *nnf1a* RNAi (109867KK), 0.14μm/sec, n=219 comets. E) Quantification graph of velocity of EB1-GFP comets in the primary protrusion of qNSCs at 6h ALH from various genotypes in (A). Control, 0.16μm/sec, n=339 comets; *ndc80* RNAi (BL33620), 0.14μm/sec, n=98 comets; *ndc80* RNAi (29337GD), 0.14μm/sec, n=65 comets; *spc25* RNAi (104213KK), 0.16μm/sec, n=214 comets; *spc25* RNAi (27923GD), 0.14μm/sec, n=109 comets; *nuf2* RNAi (BL35599), 0.13μm/sec, n=105 comets; *spc105R* RNAi (BL35466), 0.14μm/sec, n=106 comets and *spc105R* RNAi (BL36100) 0.13μm/sec, n=120 comets. F) Quantification graph of the percentage of anterograde- and retrograde-moving EB1-GFP comets in the primary protrusion of qNSCs from various genotypes in (A). Anterograde-moving comets: control, 92.7%, n=313 comets; *mis12* RNAi (105052KK), 84.5%, n=108 comets; *mis12* RNAi (BL35471), 80.5%, n=107 comets; *mis12* RNAi (BL38535), 80.8%, n=105 comets; *kmn1* RNAi (19529GD), 86.2%, n=56 comets; *kmn1* RNAi (106889KK), 77.7%, n=94 comets; *nnf1a* RNAi (18208GD), 85.4%, n=103 comets; *nnf1a* RNAi (109867KK), 86%, n=189 comets. Retrograde-moving comets: control, 7.3%, n=24 comets; *mis12* RNAi (105052KK), 15.5%, n=20 comets; *mis12* RNAi (BL35471), 19.5%, n=26 comets; *mis12* RNAi (BL38535), 19.2%, n=25 comets; *kmn1* RNAi (19529GD), 13.8%, n=9 comets; *kmn1* RNAi (106889KK), 22.3%, n=27 comets; *nnf1a* RNAi (18208GD), 14.6%, n=18 comets; *nnf1a* RNAi (109867KK), 14%, n=30 comets. G) Quantification graph of the percentage of anterograde- and retrograde-moving EB1-GFP comets in the primary protrusion of qNSCs from various genotypes in (A). Anterograde-moving comets: control, 92.7%, n=313 comets; *ndc80* RNAi (BL33620), 84.7%, n=83 comets; *ndc80* RNAi (29337GD), 81.5%, n=53 comets; *spc25* RNAi (104213KK), 85.5%, n=183 comets; *spc25* RNAi (27923GD), 81.6%, n=89 comets; *nuf2* RNAi (BL35599), 78.1%, n=82 comets; *spc105R* RNAi (BL35466), 84.9%, n=90 comets and *spc105R* RNAi (BL36100) 80.8%, n=97 comets. Retrograde-moving comets: control, 7.3%, n=24 comets; *ndc80* RNAi (BL33620), 15.3%, n=15 comets; *ndc80* RNAi (29337GD), 18.5%, n=12 comets; *spc25* RNAi (104213KK), 14.5%, n=31 comets; *spc25* RNAi (27923GD), 18.4%, n=20 comets; *nuf2* RNAi (BL35599), 21.9%, n=23 comets; *spc105R* RNAi (BL35466), 15.1%, n=16 comets and *spc105R* RNAi (BL36100) 19.2%, n=23 comets. Data information: ns, non-significant, *p<0.05, **p<0.01, ***p<0.001 and ****p<0.0001 Data are presented as mean ± SD. Statistical significance was determined by one-way ANOVA with multiple comparisons. Scale bars: 10μm.

Knockdown of kinetochore proteins also caused microtubule orientation defects, with increased percentage of retrograde EB1-GFP comet movements as compared to control (Figure 2F-G, Movie S1). To further elucidate the effect of kinetochore components on microtubule orientation, we examined the localization of a microtubule minus-end marker *Nod-β-Gal* in qNSCs. At 16h ALH, in control qNSCs, *Nod-β-Gal* was concentrated at the apical region of the primary protrusion (Figure S2C-D). By contrast, *Nod-β-Gal* was delocalized from the apical region and distributed around the cell body or protrusion initiation segment (PIS) region of the primary protrusion upon *mis12* RNAi, *nuf2* RNAi, *ndc80* RNAi, *spc25* RNAi and *spc105R* RNAi knockdown (Figure S2C-D). Therefore, these data indicate that kinetochore proteins are important for acentrosomal microtubule growth and orientation in the primary protrusion of *Drosophila* qNSCs.

Given that kinetochore complex proteins are required for microtubule growth in qNSCs, we examined whether depletion of kinetochore proteins resulted in morphological defects in the primary protrusion of qNSCs. The thickness of the primary protrusion was measured at the middle position of the protrusion marked by *grh*>CD8-GFP. The thickness of the protrusion was significantly decreased in various kinetochore knockdown lines as compared with control qNSCs (Figure S2E-F). However, the length of the primary protrusion in qNSCs in the ventral nerve cord (VNC) upon depletion of kinetochore proteins at 16h ALH was not significantly different from that of the control (Fig S2E, G), which is likely due to the relatively constant distance between the cell body of qNSCs and neuropil. Therefore, knockdown of kinetochore proteins resulted in thinning of the primary protrusion in qNSCs.

On the contrary to the knockdown phenotypes, overexpression of kinetochore proteins by *grh*-Gal4 at 6h ALH caused a significant increase in the total number of EB1 comets (1.38-fold in UAS-*Ndc80* and 1.4-fold in UAS-*Spc105*; Figure S3A-B; Movie S2) as compared to control (1-fold; Figure S3A-B; Movie S2). Overexpression of Mis12 only caused a slight but non-significant increase in total number of EB1-GFP comets (1.3-fold, Fig S3A-B). Similarly, the average velocity of EB1 comets was also significantly increased by Ndc80 or Spc105 overexpression, compared to control (Figure S3A, C; Movie S2). Furthermore, the percentage of anterograde versus retrograde moving EB1 comets, though not statistically significant was also slightly increased upon overexpression of various kinetochore proteins as compared to the control (Figure S3A, D; Movie S2).

### Kinetochore proteins are required for the regeneration of primary protrusion of qNSCs upon injury

Recently, we have demonstrated that the primary protrusion of *Drosophila* qNSCs could be regenerated upon injury, which is dependent on Golgi function ^19, 20^. Given that kinetochore proteins promote microtubule assembly in qNSCs, we tested whether kinetochore proteins are important for this regeneration. To this end, we performed laser ablation on qNSC protrusions of several kinetochore proteins knockdown by RNAi. At 12h ALH, following laser ablation at the middle region of primary protrusions, 53.9% of *mis12* RNAi 105052KK, 55.5% of *nuf2* RNAi BL35599, 44.4% of *spc25* RNAi 27923GD and 40% of *spc105R* RNAi BL35466 qNSCs in *ex vivo* larval brains failed to fully regenerate their primary protrusion within 30 minutes of imaging respectively as compared to 28.5% in the control (Figure 3A-B, Movie S3, Methods). Our data suggest that the kinetochore complex proteins are important for the regeneration of primary protrusion of *Drosophila* qNSCs upon injury.

**Figure 3.**
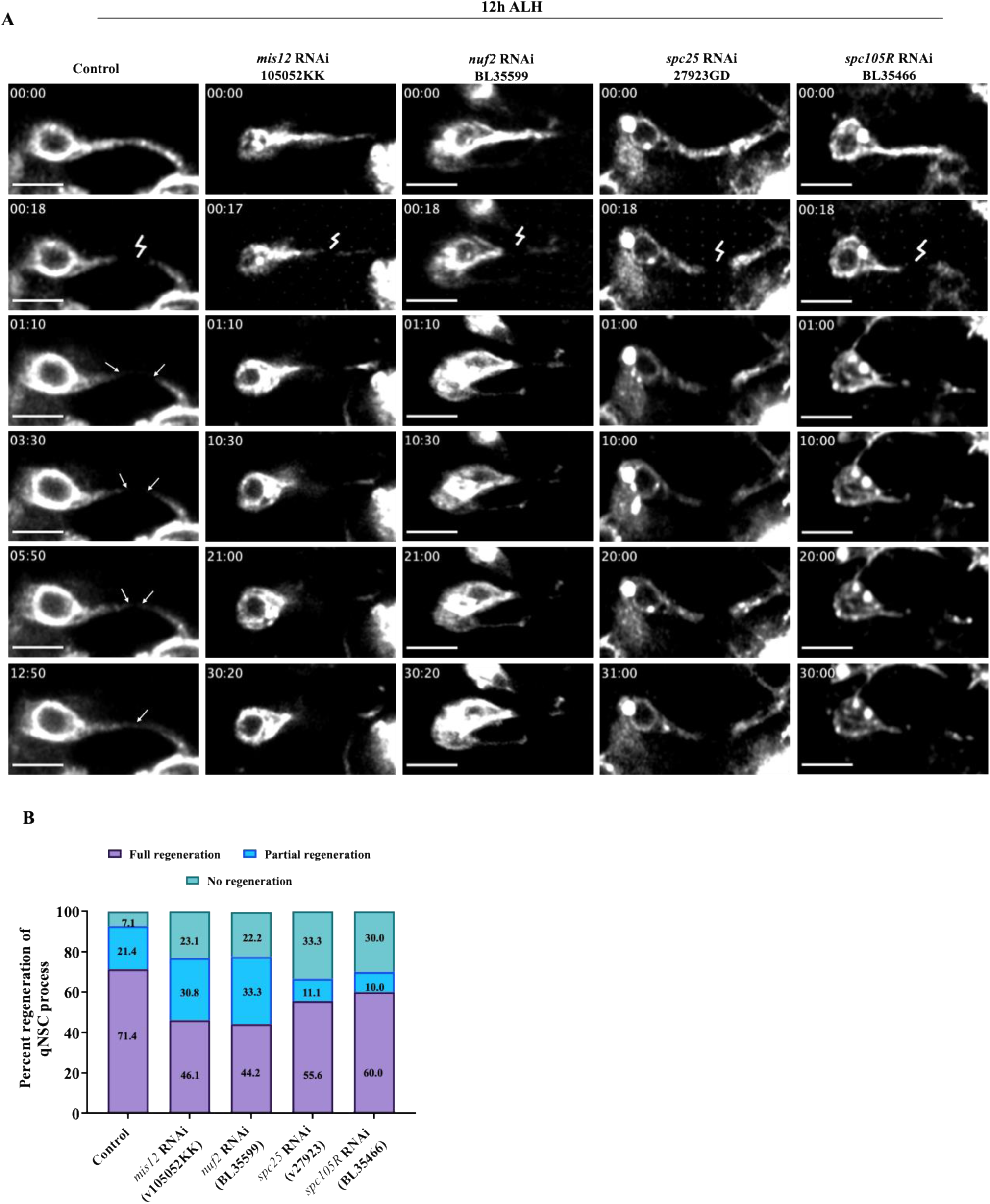
Kinetochore proteins are required for the regeneration of qNSCs. A) Time series of a quiescent NSC in *ex vivo* larval brain at 12h ALH labelled by *grh*-Gal4; UAS-mCD8-GFP in control (UAS*-β-Gal* RNAi), *mis12* RNAi (105052KK), *nuf2* RNAi (BL35599), *spc25* RNAi (27923GD) and *spc105R* RNAi (BL35466) ablated at the middle region of the protrusion (lightning strike). Arrows in control indicate regeneration of the protrusion. B) Quantification graph of percentage of regeneration of for genotypes in (A). Control, complete regeneration=71.4%, partial regeneration=21.4%, no regeneration=7.1%, n=14; *mis12* RNAi (105052KK), complete regeneration=46.1%, partial regeneration=30.8%, no regeneration=23.1%, n=13; *nuf2* RNAi (BDSC BL35599), complete regeneration=44.2%, partial regeneration=33.3%, no regeneration=22.2%, n=9; *spc25* RNAi (VDRC 27923GD), complete regeneration=55.6%, partial regeneration=11.1%, no regeneration=33.3%, n=9 and *spc105R* RNAi (BDSC BL35466), complete regeneration=60%, partial regeneration=10%, no regeneration=30%, n=10. Data information: Data are presented as mean. Scale bars: 10μm.

### Kinetochore genes have increased expression levels in active NSCs, compared with qNSCs

To determine the expression pattern of kinetochore proteins in larval brains, we analyzed a published dataset of single cell RNA-sequencing obtained from late first instar larvae at 16 h ALH ^51^. Kinetochore proteins, including Ndc80, Mis12, Spc25, Nuf2, Spc105R, Kmn1, Nnf1a and Nnf1b showed lower expression level in qNSCs, but increased notably in active NSCs (aNSCs) (Figure 4A). Proliferating markers such as Wor, CycA, CycE and Dpn, a NSC marker, that have higher expression levels in aNSCs than qNSCs, were included as positive controls, while *Rpl32*, a housekeeping gene has similar expression levels in these two cell populations (Figure 4A).

**Figure 4.**
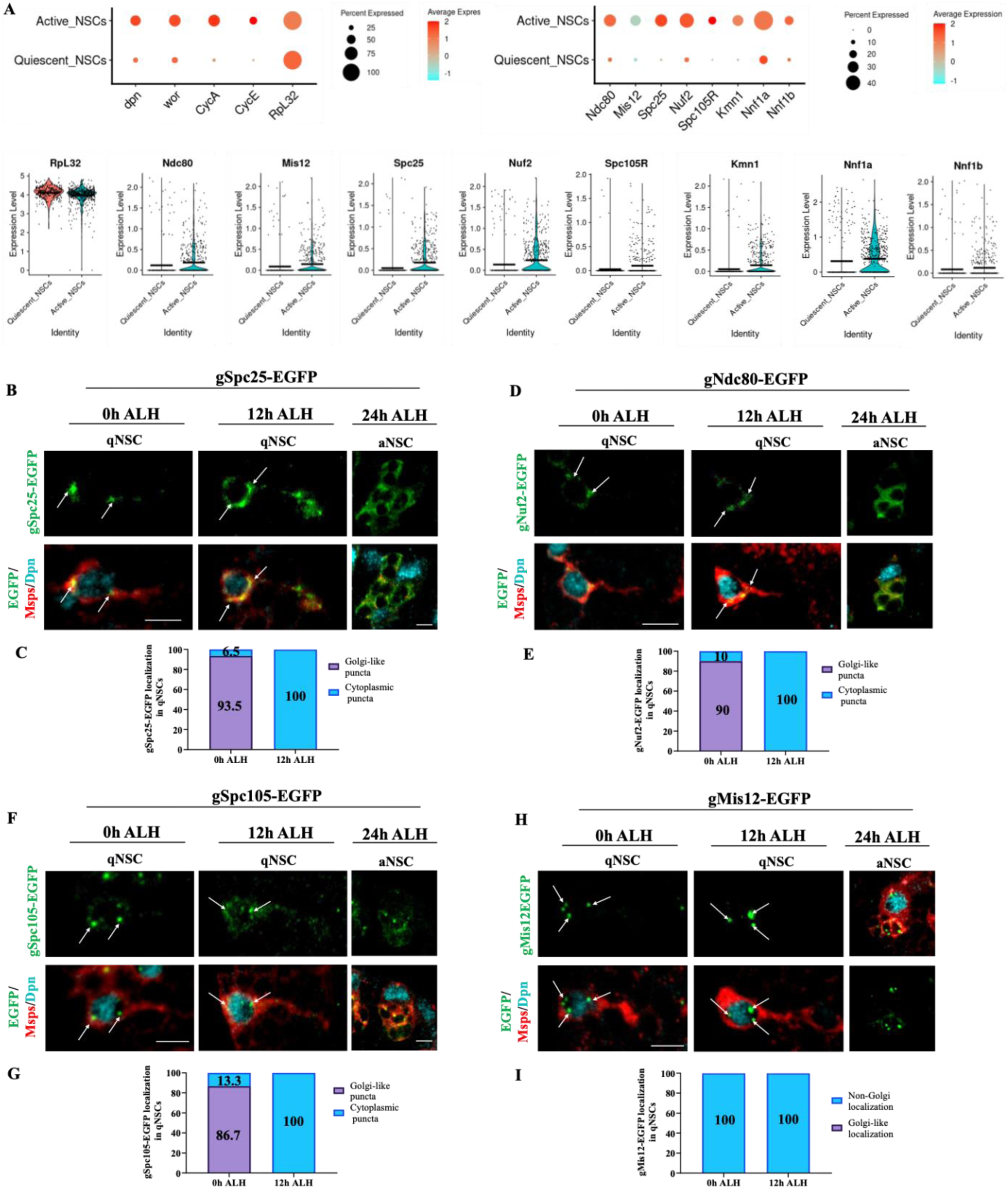
Localization of Kinetochore proteins in qNSCs. A) Analysis of kinetochore complex proteins mRNA expression in qNSCs and aNSCs from the dataset of single-cell RNA seq (Brunet et al). *RpL32* works as a house keeping gene. B) Localization of genomic Spc25-EGFP (gSpc25-EGFP) in qNSCs at 0h and 12h ALH. Localization of gSpc25-EGFP in an active NSC (aNSC) at 24h ALH. Cells are outlined by Msps and nucleus is marked by Dpn. C) Quantification graph of the percentage of qNSCs at 0h and 12hALH showing gSpc25-EGFP localization. 0h ALH, Golgi-like puncta, 93.5%; cytoplasmic puncta, 6.5%, n=31. 12h ALH, Golgi-like puncta, 0%; cytoplasmic puncta, 100%, n=30. D) Localization of genomic Ndc80-EGFP (gNuf2-EGFP) in qNSCs at 0h and 12h ALH. Localization of gNdc80-EGFP in an active NSC (aNSC) at 24h ALH. Cells are outlined by Msps and nucleus is marked by Dpn. E) Quantification graph of the percentage of qNSCs at 0h and 12hALH showing gNdc80-EGFP localization. 0h ALH, Golgi-like puncta, 90%; cytoplasmic puncta, 10%, n=30. 12h ALH, Golgi-like puncta, 0%; cytoplasmic puncta, 100%, n=34. F) Localization of genomic Spc105-EGFP (gSpc105-EGFP) in qNSCs at 0h and 12h ALH. Localization of gSpc105-EGFP in an active NSC (aNSC) at 24h ALH. Cells are outlined by Msps and nucleus is marked by Dpn. G) Quantification graph of the percentage of qNSCs at 0h and 12hALH showing gSpc105-EGFP localization. 0h ALH, Golgi-like puncta, 86.7%; cytoplasmic puncta, 13.3%, n=30. 12h ALH, Golgi-like puncta, 0%; cytoplasmic puncta, 100%, n=24. H) Localization of genomic Mis12-EGFP (gMis12-EGFP) in qNSCs at 0h and 12h ALH. Localization of gMis12-EGFP in an active NSC (aNSC) at 24h ALH. Cells are outlined by Msps and nucleus is marked by Dpn. I) Quantification graph of the percentage of qNSCs at 0h and 12hALH showing gMis12-EGFP localization. 0h ALH, Golgi-like localization, 0%; non-Golgi localization, 100%, n=34. 12h ALH, Golgi-like localization, 0%; non-Golgi localization, 100%, n=31. Golgi-like localization refers to strong genomic kinetochore localization near Golgi (either at Apical, PIS or both regions). Non-Golgi localization includes full cytoplasmic localization of EGFP or localization within the nucleus. Scale bars: 5μm for single qNSCs.

### Kinetochore proteins are distributed as puncta in close proximity to the Golgi in *Drosophila* quiescent NSCs

To assess the localization of kinetochore proteins, we made use of transgenic flies containing the genomic regions of kinetochore proteins fused to EGFP. As expected, in mitotic NSCs at 48h ALH, gNdc80-EGFP, gMis12-EGFP, gSpc25-EGFP, and gSpc105 localized to kinetochores (Figure S3E-G). Consistent with changes in mRNA levels, gSpc25-EGFP, gNdc80-EGFP, and gSpc105-EGFP signal intensity increased from 0h ALH qNSCs to at 24h ALH aNSCs (Figure 4B-G). At 48h ALH, we found that interphase NSCs showed diffused localization of genomic Ndc80 (gNdc80), gMis12 and gSpc25 around the nucleus (Figure S3E-G), similar to known Golgi distribution in various other *Drosophila* cell types.

Among these four kinetochore proteins examined, gSpc25-EGFP displayed a clear localization at both apical and PIS region in qNSCs at 0h ALH (Figure 4B-C, 93.5%, n=31), resembling the Golgi localization in these cells. Between 12h-24h ALH, gSpc25-EGFP was seen as many cytoplasmic puncta including the primary protrusion (Figure 4B-C,100% of qNSCs, n=30), likely reflecting Golgi fragmentation process when NSCs re-enter the cell cycle. Similarly, at 0h ALH gNdc80-EGFP has a Golgi-like localization pattern at apical and PIS regions in qNSCs and was re-distributed to cytoplasm along with reactivation (Figure 4D-E, 100%, n=34). Interestingly, at 0h ALH gSpc105-EGFP displayed as two major puncta close to apical and PIS regions at the border of the nucleus in qNSCs but became many diffused puncta in the nucleus by 12h ALH (Figure 4F-G). gMis12-EGFP puncta were only seen in the nucleus but most of them were located close to apical and PIS regions of qNSCs at both 0h ALH and 12h ALH (Figure 4H-I).

Our recent work demonstrated that the Golgi apparatus is localized as 1-3 distinct puncta predominantly to the protrusion initial segment (PIS) and apical region of *Drosophila* qNSCs, unlike dispersed distribution in other *Drosophila* cell types^19^. By co-labeling various kinetochore proteins with Golgi protein Arf1, we found that gSpc25-EGFP puncta colocalized with or located in close proximity to Arf1-positive Golgi in qNSCs at both 0h and 6h ALH (Figure 5A, Figure S4A). gNdc80-EGFP also had partial co-localization with Arf1, particularly at the PIS regions at both 0h and 6h ALH (Figure 5A, Figure S4A). gSpc105-EGFP did not co-localize with Arf1 but localized in the apical and PIS regions of the qNSCs close to Arf1 at 0h ALH (Figure 5A), while gMis12-EGFP appeared as large 3-4 puncta all located within the nucleus but close to apical and PIS Golgi at 6h ALH (Figure S4A). Therefore, kinetochore proteins in qNSCs are distributed as distinct puncta either co-localizing with Golgi or close to the Golgi vicinity in the cytoplasm or nucleus.

**Figure 5.** Kinetochore complex proteins localize to or near Golgi and are required for are required for Arf1 localization at Golgi in qNSCs. A) Micrograph images of qNSCs at 0h ALH in gSpc25-EGFP, gNdc80-EGFP and gSpc105-EGFP labelled with antibodies against GFP, Arf1, Msps and DNA. Line graphs show intensity curves of Arf1 (gray) with gSpc25-EGFP, gNdc80-EGFP and gSpc105-EGFP (green) along the apical region to PIS region of the qNSC. Yellow arrowheads point to Kinetochore localization at, near or away from Arf1. B) qNSC with primary protrusions labelled by mCD8-GFP at 16h ALH from control (UAS*-β-Gal* RNAi), *ndc80* RNAi (29337GD), *spc25* RNAi (27923GD), *nuf2* RNAi (BL35599), *spc105R* RNAi (BL35466), *mis12* RNAi (BL38535) and *mis12* RNAi (105052KK) driven by the *gr*h-Gal4 driver, were labelled with antibodies against Arf1, Dpn, and GFP. C) Quantification graph of average intensity of Arf1 puncta for genotypes in (B). Control, 43.1 A.U., n=40; *ndc80* RNAi (29337GD), 11.8 A.U., n=18; *spc25* RNAi (27923GD), 19 A.U., n=26; *nuf2* RNAi (BL35599), 16.4 A.U., n=29; *spc105R* RNAi (BL35466), 21.6 A.U., n=14; *mis12* RNAi (BL38535), 29.3 A.U., n=16 and *mis12* RNAi (105052KK), 27 A.U., n=20. D) Quantification graph of the percentage of Arf1 puncta present per quiescent NSC for genotypes in (B). Control= Apical + PIS=91%, Apical=4.5%, PIS=4.5%, n=22; *ndc80* RNAi (29337GD), Apical + PIS=16.7%, Apical=16.7%, PIS=25%, Absent=41.6%, n=24; *spc25* RNAi (27923GD), Apical + PIS=12%, Apical=28%, PIS=16%, Absent=44%, n=25; *nuf2* RNAi (BL35599), Apical + PIS=37.5%, Apical=25%, PIS=12.5%, Absent=25%, n=16; *spc105R* RNAi (BL35466), Apical + PIS=8%, Apical=20%, PIS=12%, Absent=60%, n=25; *mis12* RNAi (BL38535), Apical + PIS=52.2%, Apical=17.4%, PIS=13%, Absent=17.4%, n=23 and *mis12* RNAi (105052KK), Apical + PIS=34.8%, Apical=17.4%, PIS=4.3%, Absent=43.5%, n=23. Data information: Arrow heads indicate Arf1 puncta localization. Data are presented as mean ± SD. Statistical significance was determined by one-way ANOVA with multiple comparisons. *p<0.05, **p<0.01, ***p<0.001 and ****p<0.0001. Scale bars: 5μm.

### Golgi localization of Arf1 and Sec71 in qNSCs is dependent on kinetochore proteins

Next, we investigated whether kinetochore complex proteins are required for the proper localization of Golgi proteins, including Arf1 and the Arf1GEF Sec71. In all control qNSCs at 16h ALH, Arf1 is localized as 1-3 distinct puncta to the apical region distal to the protrusion and the PIS of qNSCs (Figure 5B-D; 0% defect). In contrast, Arf1 localization and intensity at PIS and the apical regions was absent or diminished in *ndc80* RNAi (83.3%), *spc25* RNAi (88%), *nuf2* RNAi (62.5%), *spc105* RNAi (92%) and *mis12* RNAi (47.8% for BL38535; 65.2% for v105052KK) qNSCs (Figure 5B-D). Similarly, Sec71 puncta intensity was absent or diminished in *ndc80* RNAi (65.2%), *spc25* RNAi (66.7%), *nuf2* RNAi (57.1%), *spc105* RNAi (63.3%) and *mis12* RNAi (42.8% for BL38535; 78.6% for v105052KK) at 16h ALH as compared to control (Figure S4B-D; 0% defect). These results suggested that Golgi localization of Arf1 and Sec71 in qNSCs requires kinetochore proteins. On the contrary, localization of gMis12, gNdc80, gSpc25 or gSpc105R was unaffected in whole brain lobes and qNSCs at 6h ALH upon treatment with Brefeldin A (BFA), a drug that disrupts Golgi localization and function by inhibiting Arf1 activation (Zhou 2014, Niu 2005), similar to DMSO-treated control (Figure S4E), suggesting that Golgi proteins are dispensable for kinetochore localization in qNSCs. Taken together, these results indicate that kinetochore proteins are important for Golgi localization of Arf1 and Sec71 in qNSCs, but not vice versa.

### Kinetochore proteins can physically associate with Arf1

To determine whether Arf1 physically associates with the kinetochore complex proteins, Flag-Arf1, EGFP-Ndc80, and Mis12-EGFP were cotransfected into S2 cells followed by coimmunoprecipitation (co-IP) experiments. Indeed, EGFP-Ndc80 was detected following immunoprecipitation of Flag-Arf1 (Figure 6A). Similarly, Flag-Arf1 was detected in the immune complex with Mis12-EGFP following immunoprecipitation (Figure 6B). This data suggests that kinetochore proteins can physically associate with Arf1.

**Figure 6.**
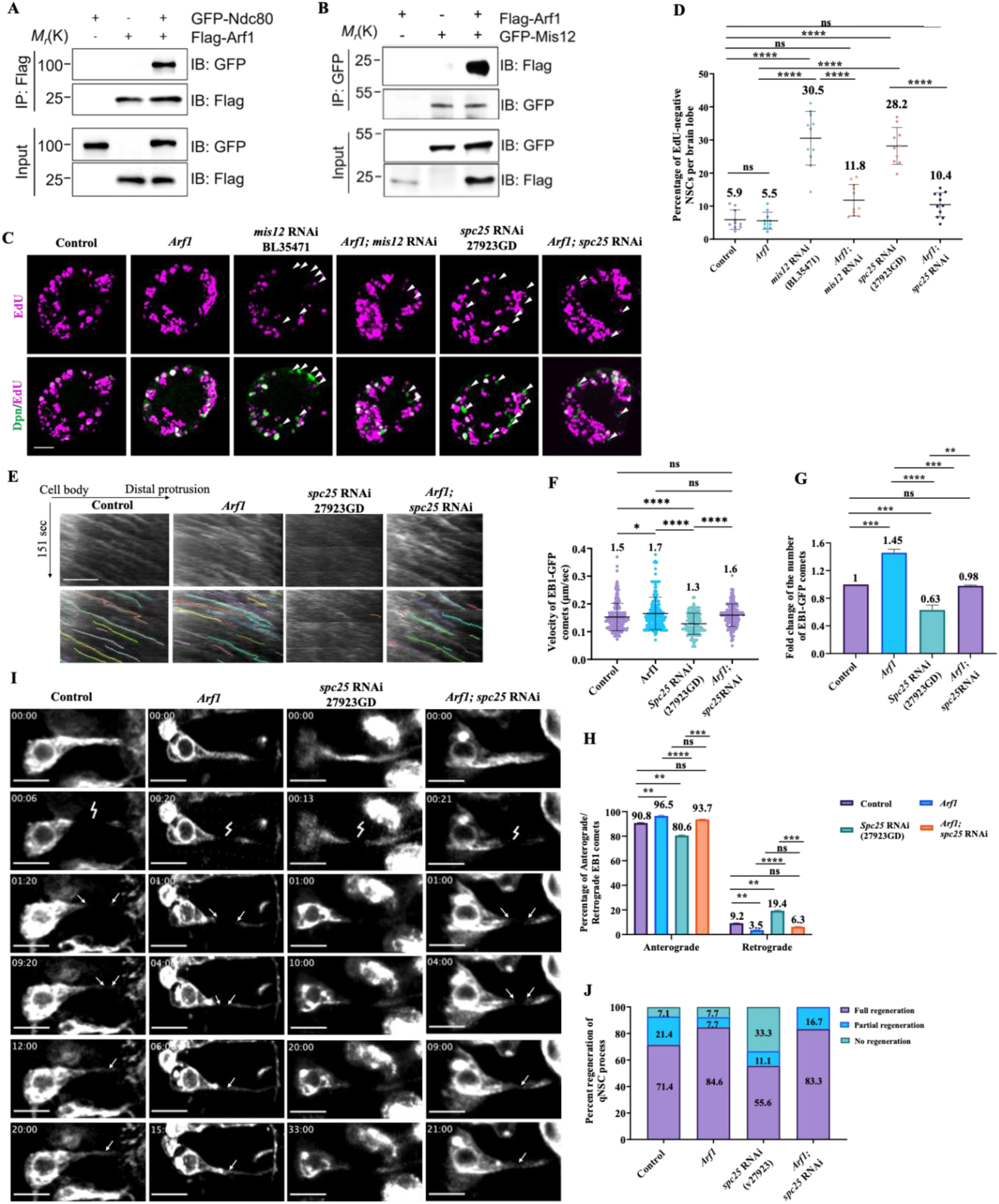
Kinetochore complex proteins physically associate with the Golgi protein Arf1 and function upstream of Arf1 to promote NSC reactivation. A-B) Co-immunoprecipitation (Co-IP) assays in S2 cells. IP was conducted with anti-Flag antibody or anti-GFP antibody and western blots were performed by using anti-Flag and anti-GFP to detect Flag-Arf1 and GFP-Ndc80 (A) or GFP-Mis12 (B), respectively. C) Larval brains at 24h ALH from control (*grh*-Gal4; UAS*-dicer2/*UAS*-β-Gal* RNAi), UAS-Arf1, *mis12* RNAi (BL35471), UAS-Arf1; *mis12* RNAi (BL35471), *spc25* RNAi (27923GD) and UAS-Arf1; *spc25* RNAi (27923GD) were analysed for EdU incorporation. NSCs were marked by Dpn. D) Quantification graph of EdU-negative NSCs per brain lobe for genotypes in (C). Control, 5.9%, n=10 BL; UAS-Arf1, 5.5%, n=10 BL; *mis12* RNAi (BL35471), 30.5%, n=11 BL; UAS-Arf1; *mis12* RNAi (BL35471), 11.8%, n=11 BL; *spc25* RNAi (27923GD), 28.2%, n=11 BL and *UAS-Arf1*; *spc25* RNAi (27923GD), 10.4%, n=10 BL. E) Kymographs of EB1-GFP comets movement in the primary protrusion of qNSCs expressing EB1-GFP under *grh*-Gal4 from the control, UAS-Arf1, *spc25* RNAi (27923GD) and UAS-Arf1; *spc25* RNAi (27923GD) at 6h ALH. The horizontal arrow indicates anterograde movement direction from cell body to the tip of the primary protrusion in qNSCs. F) Quantification graph of fold changes of number of EB1-GFP comets in the primary protrusion of qNSCs 6h ALH from various genotypes in (E). Control, 1, n=16 qNSCs, n=207 comets; UAS-Arf1, 1.45, n=12 qNSCs, n=226 comets; *spc25* RNAi (27923GD), 0.63 n=17 qNSCs, n=139 comets and UAS-Arf1; *spc25* RNAi (27923GD), 0.98, n=15 qNSCs, n=190 comets. G) Quantification graph of velocity of EB1-GFP comets in the primary protrusion of qNSCs at 6h ALH from various genotypes in (E). Control, 0.15μm/sec, n=207 comets; UAS-Arf1, 0.17μm/sec, n=226 comets; *spc25* RNAi (27923GD), 0.13μm/sec, n=139 comets and UAS-Arf1; *spc25* RNAi (27923GD), 0.16μm/sec, n=190 comets. H) Quantification graph of the percentage of anterograde- and retrograde-moving EB1-GFP comets in the primary protrusion of qNSCs from various genotypes in (E). Anterograde-moving comets: control, 90.8%, n=188 comets; UAS-Arf1, 96.5%, n=218 comets; *spc25* RNAi (27923GD), 80.6%, n=112 comets and UAS-Arf1; *spc25* RNAi (27923GD), 93.7%, n=178 comets. Retrograde-moving comets: control, 9.2%, n=19 comets; UAS-Arf1, 3.5%, n=8 comets; *spc25* RNAi (27923GD), 19.4%, n=27 comets and UAS-Arf1; *spc25* RNAi (27923GD), 6.3%, n=12 comets. I) Time series of a quiescent NSC in *ex vivo* larval brain at 12h ALH labelled by *grh*-Gal4; UAS-mCD8-GFP in control (UAS*-β-Gal* RNAi), UAS-Arf1, *spc25* RNAi (27923GD) and UAS-Arf1; *spc25* RNAi (27923GD) ablated at the middle region of the protrusion (lightning strike). Arrows indicate regeneration of the protrusion. J) Quantification graph of percentage of regeneration of for genotypes in (I). Control, complete regeneration=71.4%, partial regeneration=21.4%, no regeneration=7.1%, n=14; UAS-Arf1, complete regeneration=84.6%, partial regeneration=7.7%, no regeneration=7.7%, n=12; *spc25* RNAi (27923GD), complete regeneration=55.6%, partial regeneration=11.1%, no regeneration=33.3%, n=9 and UAS-Arf1; *spc25* RNAi (27923GD), complete regeneration=83.3%, partial regeneration=16.7%, n=6. Data information: (C-D) EdU incorporation was analyzed at 24h ALH by feeding larvae at 20h ALH with food supplemented with 0.2 mM EdU for 4h. White arrowheads point to NSCs without EdU incorporation. Data are presented as mean ± SD. Statistical significance was determined by one-way ANOVA with multiple comparisons. ns, non-significant, *p<0.05, **p<0.01, ***p<0.001 and ****p<0.0001. Scale bars: 10μm.

To further validate the association between kinetochore proteins and Arf1, we employed proximity ligation assay (PLA), a technique that enables the detection of protein-protein interactions with high specificity and sensitivity ^52^. We co-expressed various proteins tagged with EGFP or Flag in S2 cells and quantified PLA foci that indicated protein interactions (Figure S5A-C). The vast majority of S2 cells co-expressing both Flag and EGFP controls had no PLA signals, except for a small number of the cells displaying a weak PLA fluorescence signal of no more than 10 foci (Figure S5A-C). Similarly, the vast majority of cells (87.2%-96.4%) co-expressing Flag-Arf1^WT^ with control EGFP and Mis12-EGFP/EGFP-Ndc80/EGFP-Nuf2/Spc25-EGFP or Spc105-EGFP with control Flag had no PLA signal; less than 1 PLA foci per cell, were detected under each co-expression condition (Figure S5A-C). By contrast, 100% of cells co-expressing Flag-Arf1^WT^ and Mis12-EGFP displayed PLA signal (20.7 PLA foci per cell on average), of which 20.9% displayed strong signal (>30 foci), 48.8% displayed moderate signal (11-30 foci), and 30.2% displayed weak signal (1-10 foci) (Figure S5A-C). Similarly, cells co-expressing Flag-Arf1^WT^ and EGFP-Ndc80 showed an average of 28 PLA foci per cell, of which 35.5% displayed strong signal (>30 foci), 55.6% displayed moderate signal (11-30 foci), and 8.9% displayed weak signal (1-10 foci) (Figure S5A-C). Flag-Arf1^WT^ and EGFP-Nuf2 co-expressing cells displayed PLA foci wherein 6.8% had strong signal, 54.6% moderate signal, and 38.6% weak signal, with an average of 15.6 PLA foci per cell (Figure S5A-C). Cells co-expressing Flag-Arf1^WT^ and Spc25-EGFP displayed PLA signal (25.4 PLA foci per cell on average), of which 28.3% displayed strong signal, 56.5% displayed moderate signal (11-30 foci), and 15.2% displayed weak signal (Figure S5A-C). Lastly, 100% of Flag-Arf1^WT^ and Spc105-EGFP co-expressing cells showed 23.8% strong PLA foci, 47.6% moderate signal, and 28.6% displayed weak signal, with 21.9 PLA foci per cell on average (Figure S5A-C). Taken together, our data indicate that Arf1 can physically associate with kinetochore proteins Mis12, Ndc80, Nuf2, Spc25 and Spc105.

### The Kinetochore complex functions upstream of Arf1-Msps and downstream of Patronin to promote NSC reactivation

The Patronin-Arf1-Msps genetic pathway regulates NSC reactivation and regeneration via acentrosomal microtubule growth ^19, 20^. We sought to investigate the epistasis of Patronin, Arf1, and Msps along with kinetochore proteins during NSC reactivation. First, we overexpressed *arf1* in *mis12* and *spc25-*depleted brains and found a strong suppression of NSC reactivation phenotypes at 24h ALH, with the number of EdU-negative NSCs dramatically reduced to 11.8% compared with 30.5% in *mis12* RNAi control brains (Figure 6C-D). Similarly, overexpression of *arf1* in *spc25* RNAi brains also showed strong suppression of NSC reactivation phenotypes (10.4% in *spc25*–depleted brains overexpressing *arf1* compared to 28.2% in *spc25* RNAi brains (Figure 6C-D). Next, we assessed whether overexpression of the Msps in *ndc80-*depleted brains could suppress the NSC reactivation defects. At 24h ALH, only 10.4% of EdU-negative NSCs were observed in *ndc80* RNAi with *Msps* overexpression compared with 22.6% in *ndc80* RNAi brains alone (Figure S6A-B). These data suggest that Arf1 and Msps function downstream of the kinetochore complex in regulating NSC reactivation.

Interestingly, in *patr^sk1^/patr^sk8^*mutant brains overexpressing *Mis12*, the number of EdU-negative NSCs was significantly reduced to 43.3% compared with 77.9% in *patronin* mutant brains (Figure S6C-D). In contrast, overexpressing *Patronin* in *ndc80*-depleted brains did not suppress the NSC reactivation phenotypes (Figure S6E-F), suggesting that kinetochore proteins function downstream of Patronin. Taken together, these data suggest the Patronin-Kinetochore proteins-Arf1-Msps genetic pathway in promoting NSC reactivation.

To further examine the epistasis of Arf1 and Msps along with kinetochore proteins during acentrosomal microtubule growth in quiescent NSCs, we overexpressed *arf1* in *spc25-*depleted brains and found that *arf1* could significantly suppress the defects in the number, velocity and orientation of EB1-GFP comets caused by the loss of *spc25* (Figure 6E-G; Movie S4). Similarly, over-expressing *Msps* could significantly rescue the EB1-GFP number and velocity defects caused by *ndc80* loss (Figure S6G-J; Movie S5). These data suggest that the Kinetochore complex-Arf1-Msps pathway drives acentrosomal microtubule growth in qNSCs. To determine whether Arf1 functions downstream of kinetochore proteins in regulating regeneration of qNSC cellular protrusion after injury, we overexpressed *arf1* in *spc25-*depleted brains and found that *arf1* could significantly suppress the defects in regeneration of cellular protrusion caused by the loss of *spc25* (Figure 6I-J; Movie S6). Thus, kinetochore proteins likely function upstream of Arf1 in regulating regeneration of qNSC protrusion after injury.

### Human NDC80/HEC1 and SPC25 localize to or near Golgi in HEK293T cells

Given that we found that *Drosophila* kinetochore proteins could localize near or at the Golgi in qNSCs, we next tested the localization patter of human kinetochore proteins SPC25 and NDC80/HEC1 in HEK293T cells. In mitotic HEK293T cells, SPC25 and NDC80 localized to kinetochores as expected (Figure 7A, C). Interestingly, in interphase cells, SPC25 was seen as a cluster of puncta colocalized with a Golgi marker GM130 (Figure 7B). Intensity of SPC25 in both mitotic and interphase cells was significantly reduced upon knockdown of *SPC25* by siRNA (Figure S7A), suggesting the specificity of the antibodies in immunohistochemistry. NDC80 localized throughout the cytoplasm of interphase HEK293T, slightly overlapping with GM130 (Figure 7D). Therefore, similar to Golgi localization of *Drosophila* kinetochore proteins in qNSCs, some human kinetochore proteins could localize to Golgi vicinity in interphase human cells.

**Figure 7.**
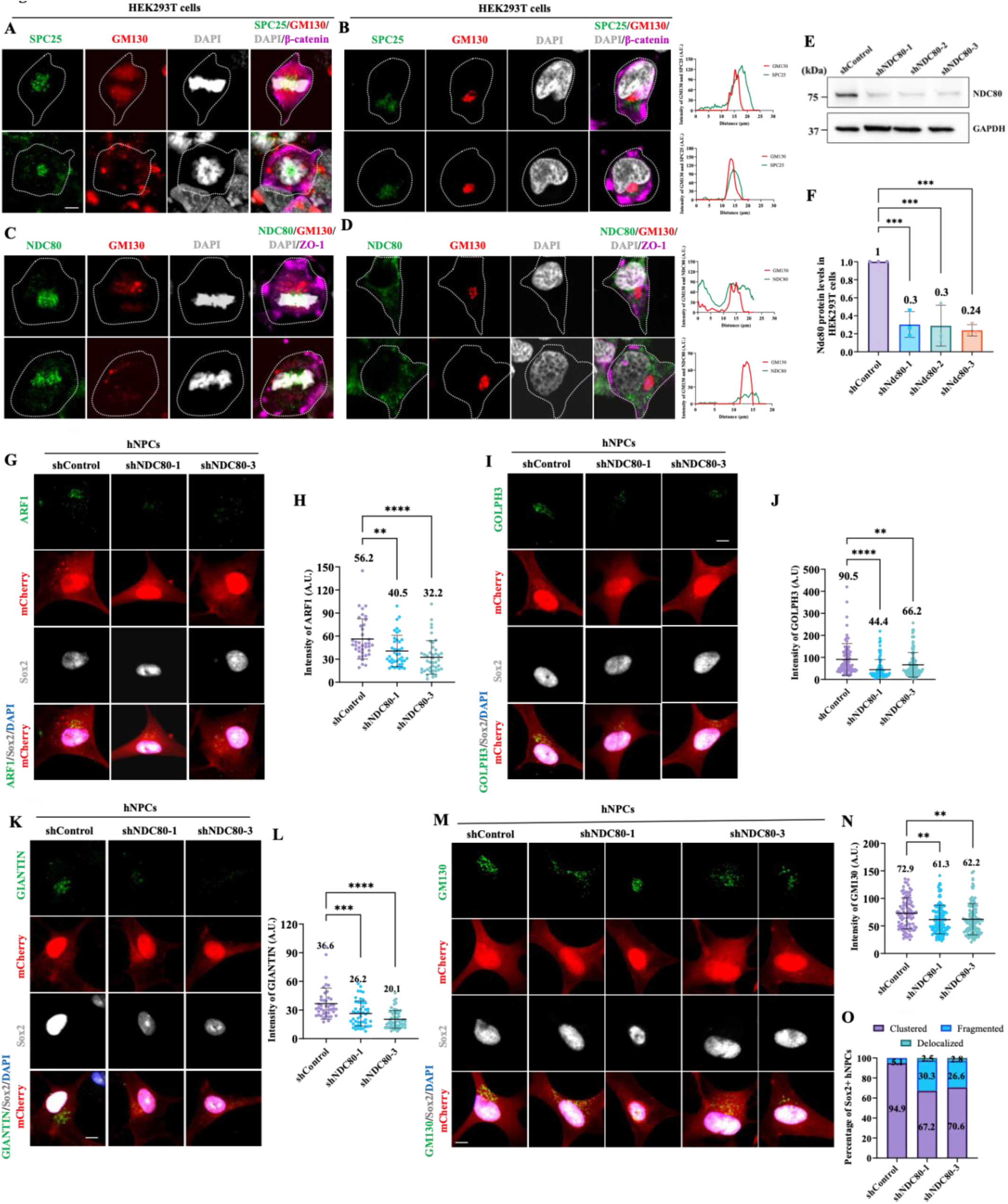
Kinetochore protein NDC80 is required for Golgi localization in hNPCs. A) Immunostaining micrographs showing human SPC25 localization in mitotic HEK293T cells. SPC25 localizes as punctate structures near the chromosomes. Cells are labelled by SPC25 (green), GM130 (red), DAPI (gray) and β-catenin (magenta, to mark cell outline). N=42. B) Immunostaining micrographs showing human SPC25 localization in interphase HEK293T cells. SPC25 localizes in the cytoplasm of the cells especially near Golgi. Cells are labelled by SPC25 (green), GM130 (red), DAPI (gray) and β-catenin (magenta, to mark cell outline). N=65 C) Immunostaining micrographs showing human NDC80 localization in mitotic HEK293T cells. NDC80 localizes as punctate structures near the chromosomes. Cells are labelled by NDC80 (green), GM130 (red), DAPI (gray) and ZO-1 (magenta, to mark cell outline). N=46. D) Immunostaining micrographs showing human NDC80 localization in interphase HEK293T cells. NDC80 localizes in the cytoplasm of the cells overlapping with GM130. Cells are labelled by NDC80 (green), GM130 (red), DAPI (gray) and ZO-1 (magenta, to mark cell outline). N=69. E) WB analysis of HEK293T cell protein extracts from control (shControl-mCherry) and three different NDC80 KDs (shNDC80-1, shNDC80-2 and shNDC80-3) with lentivirus infection in 72h culture. Blots were probed with anti-NDC80 antibody and anti-GAPDH antibody. Immunoblot showing the efficiency of NDC80 KD. F) Quantification graph representing fold change in knockdown efficiency of NDC80 normalized to the internal control GAPDH and shControl. shControl, 1, n=3; shNDC80-1, 0.3, n=3; shNDC80-2, 0.29, n=3; shNDC80-3, 0.24, n=3. G) Immunostaining micrographs showing ARF1 localization in shControl, shNDC80-1 and shNDC80-3 hNPCs 72h after transfection. Transfected cells are labelled by endogenous mCherry. hNPCs are identified by the presence of SOX2. DNA is stained by DAPI. H) Quantification graph of average intensity of ARF1 for genotypes in (G). shControl, 56.2 A.U., n=38; shNDC80-1, 40.5 A.U., n=41; shNDC80-3, 32.2 A.U., n=42. I) Immunostaining micrographs showing GOLPH3 localization shControl, shNDC80-1 and shNDC80-3hNPCs 72h after transfection. Transfected cells are labelled by endogenous mCherry. hNPCs are identified by the presence of SOX2. DNA is stained by DAPI. J) Quantification graph of average intensity of GOLPH3 for genotypes in (I). shControl, 90.5 A.U., n=79; shNDC80-1, 44.4 A.U., n=94; shNDC80-3, 66.2 A.U., n=107. K) Immunostaining micrographs showing GIANTIN localization in shControl, shNDC80-1 and shNDC80-3 hNPCs 72h after transfection. Transfected cells are labelled by endogenous mCherry. hNPCs are identified by the presence of SOX2. DNA is stained by DAPI. L) Quantification graph of average intensity of GIANTIN for genotypes in (K). shControl, 36.6 A.U., n=43; shNDC80-1, 26.2 A.U., n=49; shNDC80-3, 20.2 A.U., n=55. M) Immunostaining micrographs showing GM130 localization in shControl, shNDC80-1 and shNDC80-3 hNPCs 72h after transfection. Transfected cells are labelled by endogenous mCherry. hNPCs are identified by the presence of SOX2. DNA is stained by DAPI. N) Quantification graph of average intensity of GM130 for genotypes in (M). shControl, 72.9 A.U., n=98; shNDC80-1, 61.3 A.U., n=119; shNDC80-3, 62.2 A.U., n=109. O) Quantification graph of clustered, fragmented or delocalized GM130 for genotypes in (M). shControl, Clustered Golgi= 94.9%, Fragmented Golgi=5.1%, Delocalized Golgi=0%, n=98; shNDC80-1, Clustered Golgi= 67.2%, Fragmented Golgi=30.3%, Delocalized Golgi=2.5%, n=119; shNDC80-3, Clustered Golgi= 70.6%, Fragmented Golgi=26.6%, De-localized Golgi=2.8%, n=109. Data information: ns, non-significant, *p<0.05, **p<0.01, ***p<0.001 and ****p<0.0001 Data are presented as mean ± SD. Statistical significance was determined by one-way ANOVA with multiple comparisons. Scale bars: 10μm.

### Human Ndc80/HEC1 and MIS12 promote Golgi assembly in human interphase cells

To determine if mammalian kinetochore proteins have a conserved function in regulating Golgi localization, we knocked down NDC80/HEC1 in both HEK293T cells and human neural progenitor cells (hNPCs). All three shRNAs were effective in knocking down of human NDC80/HEC1 in HEK293T cells (Figure 7E-F). All subsequent experiments were conducted using shNDC80-1 and shNDC80-3. To determine whether human NDC80/HEC1 is required for Golgi protein localization, we examined the localization of several Golgi proteins, including ARF1 that localized to both trans- and cis-Golgi, a trans-Golgi marker GOLPH3, cis/medial-Golgi marker GIANTIN and cis-Golgi marker GM130. Since the Golgi apparatus in the mammalian cells has a distinct localization as a perinuclear cluster during interphase and is dispensed throughout the cytosol during mitosis, we focused on their localization in interphase cells. Remarkably, knockdown of *NDC80* in both hNPCs and HEK293T interphase cells resulted in a dramatic reduction in the intensity of ARF1, GOLPH3 and GIANTIN as compared to control (Figure 7G-L; Figure S7B-G). Knockdown *NDC80* did not seem to strongly affect GM130 intensity but showed a higher number of cells with dispersed, fragmented or GM130 delocalized to the nucleus as compared to shControl (Figure 7M-O; Figure S7H-J). This moderate perturbation of GM130 is likely because it is relatively more stably tethered to the cis-Golgi matrix and has reduced dynamic cycling ^53^.

Next, we tested the effect of human NDC80 overexpression, which showed a 6.4-fold increase of its protein levels compared with the control in HEK293T cells (Figure 8A-B), verifying NDC80 overexpression construct. Notably, NDC80 overexpression in hNPCs showed a significant increase in the intensity of all Golgi proteins, including ARF1, GOLPH3, GIANTIN, and GM130, as compared to the control (Figure 8C-J). Similar effect was observed in HEK293T cells upon human NDC80 overexpression (Figure S7K-M). Thus, human NDC80 has a conserved function in promoting Golgi protein localization in both hNPCs and HEK293T cells.

**Figure 8.**
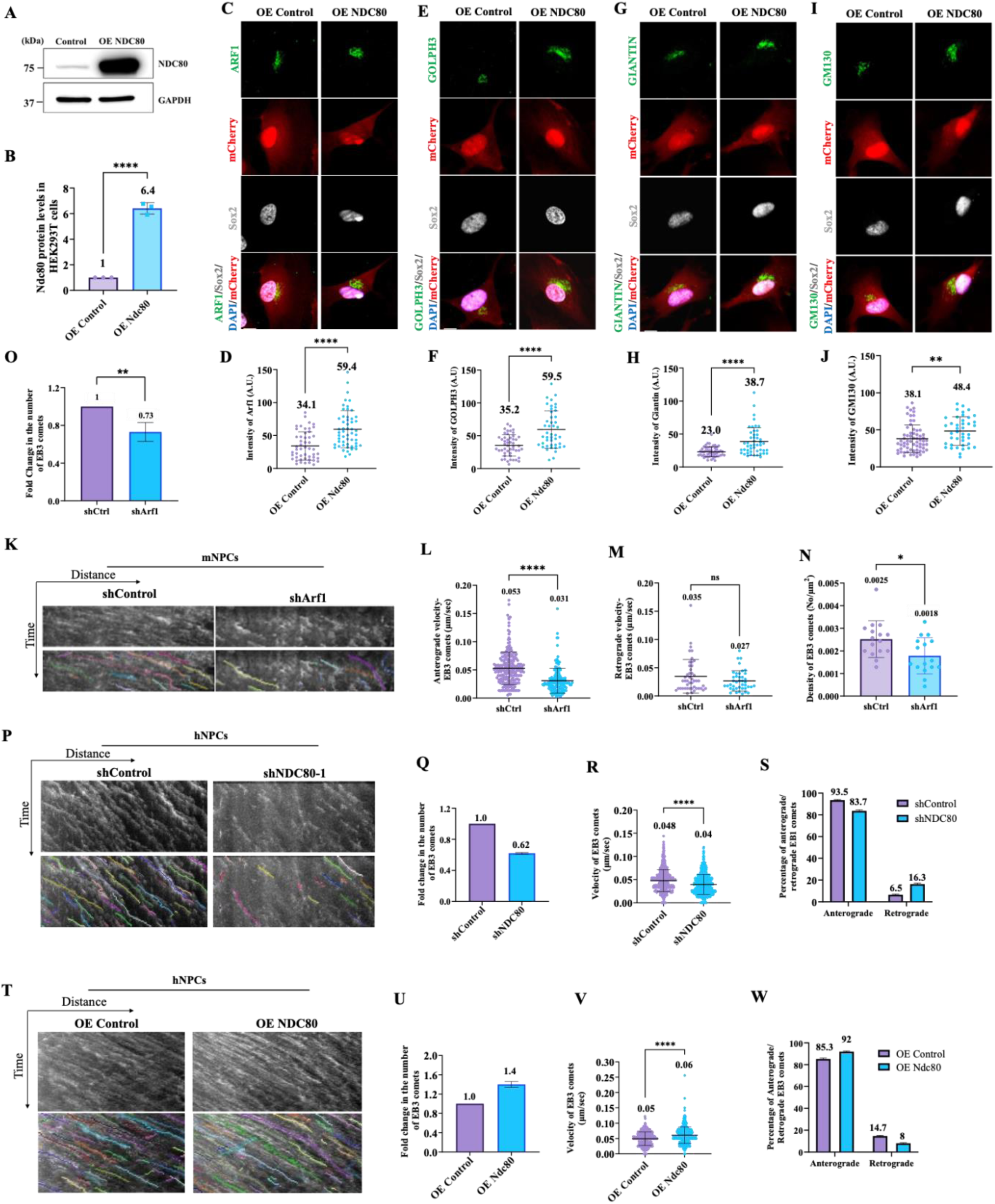
Mammalian Arf1 and Kinetochores promote microtubule growth. A) WB analysis of HEK293T cell protein extracts from control (Over expression (OE) Control-mCherry) and OE NDC80 with lentivirus infection in 72h culture. Blots were probed with anti-NDC80 antibody and anti-GAPDH antibody. Immunoblot showing the efficiency of NDC80 over expression. B) Quantification graph representing fold change in over expression efficiency of NDC80 normalized to the internal control GAPDH and OE Control. OE Control, 1, n=3; OE NDC80, 6.2, n=3. C) Immunostaining micrographs showing ARF1 localization in OE Control and OE NDC80 hNPCs 72h after transfection. Transfected cells are labelled by endogenous mCherry. hNPCs are identified by the presence of SOX2. DNA is stained by DAPI. D) Quantification graph of average intensity of ARF1 for genotypes in (C). OE Control, 34.1 A.U., n=50; OE NDC80, 59.5 A.U., n=54. E) Immunostaining micrographs showing GOLPH3 localization in OE Control and OE NDC80 hNPCs 72h after transfection. Transfected cells are labelled by endogenous mCherry. hNPCs are identified by the presence of SOX2. DNA is stained by DAPI. F) Quantification graph of average intensity of GOLPH3 for genotypes in (E). OE Control, 35.2 A.U., n=49; OE NDC80, 59.5 A.U., n=45. G) Immunostaining micrographs showing GIANTIN localization in OE Control and OE NDC80 hNPCs 72h after transfection. Transfected cells are labelled by endogenous mCherry. hNPCs are identified by the presence of SOX2. DNA is stained by DAPI. H) Quantification graph of average intensity of GIANTIN for genotypes in (G) OE Control, 23 A.U., n=53; OE NDC80, 38.7 A.U., n=48. I) Immunostaining micrographs showing GM130 localization in OE Control and OE NDC80 hNPCs 72h after transfection. Transfected cells are labelled by endogenous mCherry. hNPCs are identified by the presence of SOX2. DNA is stained by DAPI. J) Quantification graph of average intensity of GM130 for genotypes in (I). OE Control, 38.1 A.U., n=65; OE NDC80, 48.4 A.U., n=41. K) Kymographs showing the EB3 tagged with Tdtomato (Td) comets movement in mNPCs in both shControl and shArf1 groups. L and M) Quantification graphs representing the velocity of anterograde EB3 comets (shControl: 0.053 μm/s, n=16 mNPCs, n=236 comets; shArf1: 0.031 μm/s, n=16 mNPCs, n=161 comets), the velocity of retrograde EB3 comets (shControl: 0.035 μm/s, n=16 mNPCs, n=46 comets; shArf1: 0.027 μm/s, n=16 mNPCs, n=39 comets) N and O) Quantification graphs representing the total density of EB3 comets (shControl: 0.0025 No./μm2, n=16 mNPCs, n=282 comets vs. shArf1: 0.0018 No./μm2, n=16 mNPCs, n=200 comets) and the fold change in total number of EB3 comets (shControl: 1, n=16 mNPCs, n=282 comets; shArf1: 0.73, n=16 mNPCs, n=200 comets). P) Kymographs of EB3-GFP comets movement in hNPCs expressing EB3-GFP in shControl and shNDC80-1. Q) Quantification graph of fold changes of number of EB3-GFP comets in hNPCs from various genotypes in (P). Control, 1, n=25 hNPCs, n=934 comets; shNDC80-1, 0.62, n=36 hNPCs, n=838 comets. R) Quantification graph of velocity of EB3-GFP comets in comets in hNPCs from various genotypes in (P). shControl, 0.048μm/sec, n=934 comets; shNDC80-1, 0.04μm/sec, n=838 comets. S) Quantification graph of the percentage of anterograde- and retrograde-moving EB3-GFP comets in hNPCs from various genotypes in (L). Anterograde-moving comets: shControl, 93.5%, n=875 comets; shNDC80-1, 83.7%, n=700 comets. Retrograde-moving comets shControl, 6.5%, n=59 comets; shNDC80-1, 16.3%, n=138 comets. T) Kymographs of EB3-GFP comets movement in hNPCs expressing EB3-GFP in OE Control and OE NDC80. U) Quantification graph of fold changes of number of EB3-GFP comets in hNPCs from various genotypes in (K). OE Control, 1, n=14 hNPCs, n=389 comets; OE NDC80, 1.4, n=14 hNPCs, n=578 comets. V) Quantification graph of velocity of EB3-GFP comets in comets in hNPCs from various genotypes in (K). OE Control, 0.05μm/sec, n=389 comets; OE NDC80, 0.06μm/sec, n=578 comets. W) Quantification graph of the percentage of anterograde- and retrograde-moving EB3-GFP comets in hNPCs from various genotypes in (K). Anterograde-moving comets: OE Control, 85.3%, n=332 comets; OE NDC80, 92%, n=532 comets. Retrograde-moving comets OE Control, 14.7%, n=57 comets; OE NDC80, 8%, n=46 comets. Data information: ns, non-significant, *p<0.05, **p<0.01, ***p<0.001 and ****p<0.0001 Data are presented as mean ± SD. Statistical significance was determined by unpaired student’s T-test. Scale bars: 10μm.

We also knocked down another kinetochore component human MIS12 in HEK293T cells. Knockdown of MIS12 by 3 different shRNAs showed a significant reduction in protein levels (Figure S8A-B), whereas overexpression could significantly increase MIS12 protein levels by 2.9-fold (Figure S8C-D). Similar to human NDC80, knockdown of *MIS12* in HEK293T cells showed a significant reduction in the intensity of ARF1, GOLPH3, GIANTIN and GM130 as compared to control (Figure S8E-L). In contrast, MIS12 overexpression in HEK293T showed a significant increase in the intensity of all Golgi proteins, including ARF1, GOLPH3, GIANTIN, and GM130, as compared to the control (Figure S8M-T).

In mammalian cells, Golgi peri-nuclear localization requires centrosomes and centrosomal-dependent microtubules ^54^. However, knockdown of human NDC80/HEC1 or MIS12 did not affect centrosomal localization of CDK5RAP2 in interphase HEK293T cells (Figure S9A-C).

Taken together, human kinetochores promote Golgi assembly independent of centrosomes in interphase human cells.

### Mammalian Arf1 and Kinetochores promote microtubule growth

Our previous work in *Drosophila* established the Golgi protein Arf1 as a regulator of acentrosomal microtubule growth in the primary protrusion of quiescent NSCs. To understand if the function of Arf1 in regulating microtubules is conserved in mammalian cells we performed live imaging in mouse neural progenitor cells (mNPCs) to track the growing ends of microtubules by using the plus-end microtubule binding protein EB3, that is enriched in the nervous system ^55^. EB3 has been widely used to track microtubule dynamics, as it marks the microtubule growth and dynamics by binding to the plus-ends of microtubules. Remarkably, knockdown Arf1 (shArf1 marked by GFP) in mNPCs expressing EB3-tdTomato resulted in a significant reduction in velocity of anterograde moving comets (0.031 μm/s) as compared to control (Figure 8K-L and Movie S7; anterograde, 0.053 μm/s) and a slight but not significant reduction in velocity of retrograde moving comets (0.027 μm/s) as compared to control (Figure 8K, M and Movie S7; retrograde, 0.035 μm/s). Furthermore, the total density as well as fold change in total number of EB3 comets was notably reduced as compared to the control (Figure 8K, N-O; Movie S7; Density; shControl = 0.0025 No./μm2; shArf1 = 0.0018 No./μm2; Fold change in total no. of EB3 comets, shControl, 1, shArf1=0.73). Thus, our data suggests that Arf1 is required for microtubule growth in mNPCs.

To determine the role of NDC80 in Golgi-dependent microtubule growth of hNPCs, we performed live imaging using EB3-GFP. Remarkably, *NDC80* knockdown in hNPCs resulted in a dramatic reduction in the number of EB3-GFP comets compared to control (Fold change of 0.62 in shNDC80-1 compared to 1 in control; Figure 8P-Q, Movie S8). *NDC80* knockdown also caused a significant decrease in the overall velocity of both anterograde and retrograde EB3 comets as compared to control (0.048μm/s in shControl vs 0.040μm/s in shNDC80-1; Figure 8P, R, Movie S8). Furthermore, the percentage of retrograde comets in *NDC80* knockdown was notably increased as compared to the control (16.3% in shNDC80-1 as compared to 6.5% in shControl; Figure 8P, S, Movie S8). In the contrary, overexpression of NDC80 (OE NDC80) significantly increased the number of EB3 comets by 1.4-fold as compared to control (Figure 8T-U; Movie S9). Furthermore, NDC80 overexpression caused a significant increase in overall velocity of EB3 comets as compared to control (0.05μm/s in OE Control vs 0.06μm/s in OE NDC80, Figure 8T, V; Movie S9). Human NDC80 overexpression also caused a substantial increase in the percentage of anterograde moving EB3 comets as compared to OE control in hNPCs (Figure 8T, W; Movie S9).

Taken together, human NDC80 has a conserved function in promoting Golgi protein localization and microtubule growth in both hNPCs and HEK293T cells.

## Discussion

### Non-canonical functions of kinetochore proteins in non-mitotic cells

Kinetochore proteins are best known for their essential role in chromosome segregation during mitosis via mediating the attachment of spindle microtubules to centromeric chromatin. In the present study, we uncover an unexpected non-mitotic role for the KMN network in regulating Golgi organization. Specifically, we show that components of the *Drosophila* KMN network regulate Golgi organization, Golgi-dependent acentrosomal microtubule assembly, and regeneration of quiescent NSCs after injury. These findings extend a growing body of evidence indicating that kinetochore proteins can perform non-canonical functions outside of mitosis.

Several kinetochore components have recently been implicated in cytoskeletal organization and membrane trafficking during interphase. For example, the checkpoint protein Mad1 has been shown to localize to the Golgi apparatus and regulate protein secretion and cell adhesion independently of its spindle assembly checkpoint function ^56^. Similarly, the kinesin-like kinetochore protein CENP-E has been reported to regulate centrosome stability and spindle orientation ^57^. In addition, the RZZ complex component ZW10 participates in membrane trafficking between the endoplasmic reticulum and Golgi apparatus, suggesting that kinetochore-associated proteins may play broader roles in intracellular membrane organization ^58, 59^. Given that the NDC80 complex is capable of forming cooperative arrays along microtubules that stabilize dynamic plus ends ^37, 39, 60^, it is conceivable that similar microtubule-binding properties could be repurposed in non-mitotic cells to regulate microtubule assembly outside of kinetochores. Our results demonstrate that human NDC80/HEC1 and MIS12 promote Golgi organization and Golgi-dependent microtubule growth dynamics in human interphase cells. Thus, our findings suggest that the microtubule-binding module of the KMN network may represent a versatile molecular platform that can function both in chromosome segregation during mitotic and in microtubule organization in non-mitotic cells.

In many differentiated or quiescent cells, microtubule arrays are organized independently of centrosomes through non-centrosomal microtubule organizing centers (ncMTOCs), including the Golgi apparatus, nuclear envelope, and cortical sites ^61, 62^ These acentrosomal microtubule networks are particularly important in neurons and neural progenitors, where they regulate cell polarity, intracellular trafficking, and morphological remodeling. Interestingly, the effect of kinetochore protein depletion on microtubule dynamics appears to vary between cell types. In *Drosophila* sensory neurons, depletion of kinetochore proteins results in increased microtubule plus-end dynamics in dendrites but not axons ^46^. In contrast, *Caenorhabditis elegans* kinetochore proteins Cenp-C, Ndc80, and Nuf2 localize post-mitotically to dendritic structures in amphid neurons, where they regulate dendrite extension and stabilize microtubules through direct microtubule interactions ^44^. Our data reveal that the KMN network plays a previously unrecognized role in promoting Golgi-dependent acentrosomal microtubule growth in *Drosophila* quiescent NSCs.

### Kinetochore proteins regulate Golgi organization in neural stem cells

We previously demonstrated that the Golgi apparatus functions as a non-centrosomal microtubule organizing center in *Drosophila* quiescent NSCs and that the small GTPase Arf1 and its guanine nucleotide exchange factor Sec71 regulate Golgi-dependent microtubule nucleation^19^. In the present study, we identify kinetochore proteins as novel regulators of Golgi organization in both *Drosophila* and mammalian cells. Several kinetochore proteins are enriched near the perinuclear Golgi region in quiescent NSCs and are required for proper Golgi positioning. One possible mechanism is that kinetochore proteins stabilize Golgi positioning by tethering Golgi membranes to microtubules, thereby maintaining the perinuclear Golgi architecture typical of many interphase cells. Alternatively, kinetochore proteins may regulate Golgi-associated microtubule nucleation by recruiting microtubule polymerases such as Msps/XMAP215 or coordinating microtubule minus-end stabilization through Patronin/CAMSAP proteins. Patronin/CAMSAP proteins are well-established stabilizers of microtubule minus ends and play key roles in generating non-centrosomal microtubule arrays across many cell types ^63, 64^. Consistent with this model, our epistasis analysis suggests that the KMN network acts downstream of Patronin and upstream of Arf1 and Msps to regulate acentrosomal microtubule growth. The Patronin–KMN–Arf1–Msps pathway identified here therefore provides a potential mechanism linking Golgi positioning to microtubule assembly during NSC reactivation. Given that Golgi is a potent microtubule-organizing organelle in in mammalian interphase cells ^65^, it is of great interest to investigate whether Kinetochore-Golgi regulation is widely utilized in organizing microtubule in mammalian interphase cells.

### Kinetochore proteins in neural stem cell proliferation, regeneration, and brain development

Recent work has demonstrated the importance of Kinetochore proteins in neuronal regeneration. In *Drosophila* sensory neurons, multiple kinetochore components, including members of the KMN network, are upregulated following injury and are required for efficient dendritic regeneration, with comparatively minor effects on axonal regrowth ^46^. Our previous work has shown that microtubule regulators such as Arf1, Sec71, Patronin and Msps are required for the regeneration of *Drosophila* quiescent NSCs upon injury, highlighting the significance of microtubule growth in the regeneration process. In this study, we demonstrate that Kinetochore proteins play an important role in the regeneration of *Drosophila* quiescent NSCs, likely through promoting acentrosomal microtubule growth, thereby promoting a stable cytoskeletal architecture necessary for regenerative growth.

Mutations in kinetochore components have been linked to neurodevelopmental disorders in humans. Variants in the kinetochore scaffold protein KNL1 (also known as CASC5 or hSpc105), NUF2, and a kinetochore-associated centromeric protein CENP-E have been identified in autosomal recessive primary microcephaly and related neurodevelopmental abnormalities ^49, 66–69^. These clinical observations were thought to be caused by their mitotic function for neural progenitor proliferation. Our findings on kinetochore’s function on Golgi assembly in human interphase cells including human neural progenitor cells suggest that Golgi dysfunction might additionally contribute to kinetochore variant-associated microcephaly.

Together, our findings identify a previously unrecognized role for kinetochores in regulating Golgi organization, acentrosomal microtubule assembly, and neural stem cell reactivation. These results reveal a conserved molecular pathway linking kinetochore proteins with Golgi-associated regulators of microtubule growth Arf1, and Msps. Our findings on mammalian kinetochores suggests that kinetochores may represent a general mechanism for Golgi organization and Golgi-dependent functions in mammalian interphase cells.

## Material and methods

### Fly stocks and genetics

Fly stocks and genetic crosses were raised at 25°C unless otherwise stated. Fly stocks were kept in vials or bottles containing standard fly food (0.8% *Drosophila* agar, 5.8% Cornmeal, 5.1% Dextrose, and 2.4% Brewer’s yeast). The following fly strains were used in this study: *grh*-Gal4 (A. Brand), UASt-*ArfI^WT^* (F. Yu), UASt-*Msps^FL^*, *patronin^sk1^* (F. Yu), *patronin^sk8^* (F. Yu), UAS-Venus-*Patronin* (F. Yu) and UASp-*EBI*-GFP (F. Yu).

The following stocks were obtained from Bloomington *Drosophila* Stock Center (BDSC): *nuf2^SH20^*^40^ (BDSC#29510)*, nuf2^EX50^* (BDSC#91728; generated by the imprecise excision of P(lacW)Nuf2SH2276 which results in an 644bp deletion extending from 3bp upstream of ATG to 641bp downstream of ATG within the second exon with an 11bp of residual P element sequence remaining between the breaks), UAS-*mis12* RNAi (BDSC#35471), UAS-*mis12* RNAi (BDSC#38535), UAS-*ndc80* RNAi (BDSC#33620), UAS-*nuf2* RNAi (BDSC#35599), UAS-*spc105R* RNAi (BDSC#35466), UAS-*spc105R* RNAi (BDSC#36100), gMis12-EGFP (BDSC#91740), UAS-*Mis12*-EGFP (BDSC#91743), gNdc80-EGFP (BDSC#91724), UAS-*Ndc80*-EGFP (BDSC#91726), gSpc25-EGFP (BDSC#91736), UAS-*Spc25*-EGFP (BDSC#91738), gNuf2-EGFP (BDSC#91729), UAS-*Nuf2*-EGFP (BDSC#91731), gSpc105R-EGFP (BDSC#91746), UAS-*Spc105R*-EGFP (BDSC#91750), UAS-*Nnf1a*-EGFP (BDSC#91752) and UAS-*Nnf1b*-EGFP (BDSC#91754). The following stocks were obtained from Vienna *Drosophila* Resource Center (VDRC): UAS-*msps* RNAi (105052KK), UAS-*kmn1* RNAi (19529GD), UAS-*kmn1* RNAi (106889KK), UAS-*nnf1a* RNAi (18208GD), UAS-*nnf1a* RNAi (109867KK), UAS-*ndc80* RNAi (29337GD), UAS-*spc25* RNAi (27923GD) and UAS-*spc25* RNAi (104213KK). UAS- *β-Gal* RNAi (BDSC#50680) is often used as a control UAS element to balance the total number of UAS elements in each genotype. Various RNAi knockdown or overexpression constructs were induced using *grh*-Gal4 unless otherwise stated.

All experiments were carried out at 25°C, except for RNAi knockdown or overexpression studies that were performed at 29°C, unless otherwise indicated.

### EdU (5-ethynyl-2’-deoxyuridine) incorporation assay

Larvae of various genotypes were fed with food supplemented with 0.2 mM EdU from ClickiT® EdU Imaging Kits (Invitrogen) for 4 h. The larval brains were dissected in PBS and fixed with 4% EM-grade formaldehyde in PBS for 22 min, followed by washing thrice (each wash for 10 min) with 0.3% PBST, and blocked with 3% BSA in PBST for 30 min. The incorporated EdU was detected by Alexa Fluor azide, according to the Click-iT EdU protocol (Invitrogen). The brains were rinsed twice and subjected to standard immunohistochemistry.

### Immunohistochemistry

*Drosophila* larvae were dissected in PBS, and the larval brains were fixed in 4% EM-grade formaldehyde in PBT (PBS + 0.3% Triton-100) for 22 min. The samples were processed for immunostaining as previously described ^17^. For α-tubulin immunohistochemistry, the larvae were dissected in Shield and Sang M3 medium (Sigma-Aldrich) supplemented with 10% FBS, followed by fixation in 10% formaldehyde in Testis buffer (183 mM KCl, 47 mM NaCl, 10 mM Tris-HCl, and 1 mM EDTA, pH 6.8) supplemented with 0.01% Triton X-100). The fixed brains were washed once in PBS and twice in 0.1% Triton X-100 in PBS. Images were taken using LSM710 confocal microscope system (Axio Observer Z1; ZEISS), fitted with a PlanApochromat 40×/1.3 NA oil differential interference contrast objective, and brightness and contrast were adjusted by Photoshop CS6.

For hNPCs and HEK 293T cells, the cells were grown on coverslips and fixed with 4% paraformaldehyde for 15 min at RT. Subsequently, the cells were washed with PBS twice and stored prior to staining. Cells and brain sections were consequently washed with TBS and blocked with 5% normal donkey serum in TBS with 0.1% Triton X (TBST). Respective primary antibodies were prepared with the blocking solution and incubated for 2h at RT. Subsequently, the cells were washed with TBST and proceeded with secondary fluorophore antibodies incubation for 1h at RT. The cells and tissues were washed and mounted for imaging.

The primary antibodies used in this paper were guinea pig anti-Dpn (1:1,000), rabbit anti-Dpn (1:200), mouse anti-Mira (1:50, F. Matsuzaki), rabbit anti-GFP (1:3,000; F. Yu), mouse anti-GFP (1:3000; F. Yu), rabbit anti-Msps (1:1,000, J. Raff), mouse anti-β-galactosidase (1:1,000, Promega, Cat#: Z3781), mouse anti-Sec71 (1:100, F. Yu), guinea pig anti-Arf1 (1:200, F. Yu), rabbit anti-Flag (1:1,000; Sigma-Aldrich), mouse anti-α-tubulin (1:200, Sigma, Cat#: T6199), guinea pig anti-Baz (1:500; A. Wodarz), rabbit anti-PH3 (1:200, Sigma, Cat#: 06-570), rabbit anti-GOLPH3 (1:500; Abcam ab98203), rat anti-GM130 (Biolegend, Cat#:937001), rabbit anti-GM130 (Novus. Bio, Cat#: NBP2-53420), sheep anti-GM130 (R&D, Cat#:AF8199), mouse anti-GIANTIN (Abcam, ab37266), rabbit anti-ARF1 (Proteintech, Cat#:16790-1-AP), rabbit anti-HEC1/NDC80 (Abcam, ab3613), rabbit anti-hSPC25 (Proteintech, Cat#: 26474-1-AP) and anti-rabbit MIS12 (Abcam, Cat#:ab70843). The secondary antibodies used were conjugated with Alexa Fluor 488, 555 or 647 (Jackson laboratory).

### Laser ablation of qNSCs

Larval brains of various genotypes expressing UAS-mCD8-GFP under *grh*-Gal4 at various time points were dissected in Shield and Sang M3 insect medium (Sigma-Aldrich) supplemented with 10% FBS. The *ex vivo* larval brain explant culture was supplied with fat body from wild-type third instar and live imaging of the larval brains were performed with a Nikon A1R MP confocal microscope with an Apo 40× WI λ S DIC N2, N.A 1.25 objective. The laser used was 355 nm, 300 ps pulse duration, 1 kHz repetition rate, PowerChip PNV-0150-100, teem photonics. qNSCs with protrusions attached to the neuropil were chosen and imaged for about 10-30 seconds (1sec/frame) before ablation. qNSCs were hit by 355nm UV picolaser at 100nW power for 1-2secs to cause injury ^70^. After injury, qNSCs were imaged again for 10-30 seconds followed by time-lapse imaging for at least 15 minutes (1min/frame, 7-10 z-stacks with 0.5-0.8- µm intervals). The movies and images were made and analysed with NIH ImageJ software.

### Tracking of EB1-GFP or EB3-GFP comets

Larval brains of various genotypes expressing EB1-GFP under *grh*-Gal4 at various time points were dissected in Shield and Sang M3 insect medium (Sigma-Aldrich) supplemented with 10% FBS. The larval brain explant culture was supplied with the fat body from wild-type third instar and live imaging of the larval brains were performed with LSM710 confocal microscope system using 40X Oil lens and Zoom factor 6. The brains were imaged for 151 seconds, with 83 frames acquired for each movie and the images were analysed with NIH ImageJ software. The velocity of the EB1-GFP comets were calculated and kymographs were generated using KymoButler ^71^. The general cut-off for selected tracks is as follows; Sensitivity threshold for track detection -0.2, Minimum size of detected objects (pixels) -3,

Minimum number of consecutive frames per track -3, pixel size in µm (Optional) -1frame rate in seconds (Optional). 1 by default. -1. The starting points of the tracks are at the base of the PIS region where the protrusion is extended from the cell body.

mNPCs expressing EB3-GFP or EB3-TdTomato were subjected to live-cell imaging using a super-resolution SDC-SIM equipped with a Plan-Apo objective (100× 1.45-NA). The amount and velocity of the EB3-GFP comets were calculated, and kymographs were generated using KymoButler ^71^. A cell was imaged for 3 to 5 min with a 5s time interval for each movie, and videos were generated with NIH ImageJ software.

For EB3-GFP tracking in hNPCS, hNPCs expressing EB3-GFP were subjected to live-cell imaging using LSM710 confocal microscope system using 40X Oil lens and Zoom factor 2. The amount and velocity of the EB3-GFP comets were calculated, and kymographs were generated using KymoButler ^71^. A cell was imaged for 3 to 5 min without time interval for each movie, and videos were generated with NIH ImageJ software.

### Single cell RNA-seq data analysis

Raw data was downloaded from GSE134722 ^51^ and processed by Seurat 4.0. The raw data was firstly analyzed according to the methods in ^51^. Sub-clustering was then performed on the neural stem cells cluster. Quiescent and active neural stem cells were annotated by the expression of proliferating markers: Dpn, Wor, CycA, CycE and RpL32.

### Plasmids and DNA constructs

Plasmids used in this study; Flag-Arf1 (Gift from Fengwei Yu), EGFP-Ndc80, EGFP-Nuf2, Spc25-EGFP, Mis12-EGFP and Spc105-EGFP (Gift from Christian F. Lehner).

Three shRNAs targeting different regions of human NDC80/HEC1 (shNDC80-1, shNDC80-2 and shNDC80-3), human MIS12 (shMIS12-1, shMIS12-2 and sh MIS12-3) and 1 control shRNA with scrambled sequence were designed using the lentivirus shRNA knockdown vector system with Human cytomegalovirus immediate early enhancer/promoter (CMV promoter) driving the transcription of viral RNA in packaging cells and human U6 promoter driving the expression of the shRNA (VectorBuilder). Full length human NDC80, MIS12 and 1 control over expression vector were designed using the CMV promoter and Human eukaryotic translation elongation factor 1 α1 promoter (EF1A promoter) for over expression experiments (VectorBuilder).

pLV[shRNA]-mCherry-U6>Scramble_shRNA Target Sequence: CCTAAGGTTAAGTCGCCCTCG
pLV[shRNA]-mCherry-U6>hNDC80[shRNA#1]. Target sequence: TGAATTGCAGCAGACTATTAA
pLV[shRNA]-mCherry-U6>hNDC80[shRNA#2] Target sequence: GCCATTCTTGACCAGAAATTA
pLV[shRNA]-mCherry-U6>hNDC80[shRNA#3]. Target sequence: GCAGACATTGAGCGAATAAAT
pLV[Exp]-mCherry-EF1A>hNDC80 ORF: hNDC80[NM_006101.3]
pLV[shRNA]-mCherry-U6>hMIS12[shRNA#1]. Target sequence: CAGGCCGTTGAACAGGTTATT
pLV[shRNA]-mCherry-U6>hMIS12[shRNA#2]. Target sequence: GCCCAGTGCAGATTCGCAAAT
pLV[shRNA]-mCherry-U6>hMIS12[shRNA#3]. Target sequence: TTGTTCAGGCCAAACTCAAAC
pLV[Exp]-mCherry-EF1A>hMIS12 ORF: hMIS12[NM_001258218.2]

### Cell lines, transfection and Co-immunoprecipitation

*Drosophila* S2 cells (CVCL_Z232) originally from William Chia’s laboratory (with a non-authenticated identity but have been used in the laboratory for the past 10 years) were cultured in Express Five serum-free medium (Gibco) supplemented with 2 mM Glutamine (Thermo Fisher Scientific). The S2 cell culture used in this study is free of mycoplasma contamination, inferred by the absence of small speckles of DAPI staining outside of the cell nucleus. For transient expression of proteins, S2 cells were transfected using Effectene Transfection Reagent (QIAGEN) according to the manufacturer’s protocol. S2 cells were used for Co-IP and PLA assays.

S2 cells were transfected with the indicated plasmids and cultured for 48 h; collected and lysed in NP-40 lysis buffer along with protease inhibitors (Roche) for 30 min on a rotor at 4 °C. Immunoprecipitation was performed using mouse anti-Flag antibody (1:200, Sigma, F1804-1MG) and mouse anti-GFP (1:200; F. Yu), and protein G Sepharose beads according to manufacturer’s instruction. The samples were separated by SDS–PAGE and analyzed by Western blotting. Blots were probed with the following antibodies: mouse anti-Flag antibody (1:2000, Sigma, F1804-1MG) and mouse anti-GFP (1:3000; F. Yu). The secondary antibodies used were conjugated with HRP (Horseradish Peroxidase).

### Mouse neural progenitor cells (mNPCs) culture

Mouse embryos were harvested at E14, and the dorsolateral cortex was dissected and enzymatically triturated to isolate NPCs ^72^. NPCs were suspension-cultured in Costar 6-well Clear Flat Bottom Ultra-Low Attachment Multiple Well Plates (Corning) in proliferation medium (NeuroCult Proliferation Kit (Mouse & Rat), STEMCELL) containing human EGF (10 ng/ml), human FGF2 (10 ng/ml) (Invitrogen, Carlsbad, CA), N2 supplement (1%) (GIBCO), penicillin (100 U/ml), streptomycin (100 mg/ml), and L-glutamine (2 mM) for 7 days and were allowed to proliferate to form neurospheres. DIV 7 neurospheres were dissociated into single cells using accutase, yielding 5 to 6 × 106 cells per 6-well plate. Twenty-four hours lentivirus (pPurGreen) infection, NPCs were cultured as monolayer in differentiation medium containing B27 (2%) in Neurobasal medium and were maintained for 5 to 6 days.

### HEK293T culture and lentiviruses package

Clontech’s HEK 293T cell line was cultured in D-MEM high glucose medium (Invitrogen), containing 4.5 g/L D-glucose, and 4 mM L-glutamine. For packaging viral vector, high titers of engineered lentiviruses were produced by cotransfection of lentiviral vectors, VSV-G and psPAX2 into HEK293T cells followed by ultracentrifugation of viral supernatant as previously described (Ma et al, 2015).

### Human neural progenitor cells (hNPCs) culture

H9 human embryonic stem cells (hESC) were maintained on Matrigel coated dishes in mTeSR™ Plus medium (STEMCELL Technologies). Generation of hESC-derived cerebral organoids was done using STEMdiff™ Cerebral Organoid Kit (Catalog #08570, STEMCELL Technologies). H9 human organoids were grown till day 20 and enzymatically triturated to isolate hNPCs. hNPCs were suspension-cultured in Matrigel coated Costar 6-well Clear Flat Bottom Multiple Well Plates (Corning) to form neurospheres. hNPCs were grown in proliferation media (NeuroCult NS-A Basal Medium (Stem Cell STEMCELL Technologies, Human; 05750; 450 mL) and NeuroCult NS-A Proliferation Supplement (STEMCELL Technologies, Human; 05753; 50 mL)) which are mixed to prepare Complete hNSC Proliferation Medium along with heparin (2 μg/mL; Sigma), epidermal growth factor (EGF, 20 ng/mL; invitrogen), basic fibroblast growth factor (bFGF, 20 ng/mL; invitrogen), penicillin (100 U/ml) and streptomycin (100 mg/ml). hNPCs were allowed to proliferate to yield 1 × 10^6^ cells per 6-well plate. For knockdown or overexpression assays hNPC cells were seeded onto 24-well plate with 60 mm coverslips coated with matrigel, at a density of 1.5 × 10^4^ cells/coverslip. Twenty-four hours after seeding, hNPCs were infected with lentivirus after which cells were fixed and stained 72h post transfection.

### Small interfering RNA (siRNA) transfection

siRNA against human SPC25 was obtained from Santa Cruz Biotechnology (SCBT). hSPC25 (Cat#:sc-76554-V) SPC25 shRNA(h) contains lentiviral particles that are concentrated, transduction-ready containing 3 target-specific constructs that encode 19-25 nt (plus hairpin) shRNA designed to knock down gene expression. Control siRNA (Cat#: sc-37007) consists of a scrambled sequence that will not lead to the specific degradation of any cellular message. 50nM of siControl and siSPC25 were transfected into HEK293T cells using TransIT-X2® Dynamic Delivery System (Mirus Bio). Cells were then fixed and stained 48-72h post transfection.

### Generation of transgenic flies

UAS-NYFP-Myc-*Arf1* transgenic flies were generated by P-element-mediated transformation (BestGene Inc.). BDSC 8622 [yw; P(CaryP)attP2] was used as the injection stock for site-specific insertion of UAS-NYFP-Myc-*Arf1* into chromosomal location 68A4 (BestGene Inc.). pUAST-attB*-Spc25* RNAi resistant flies were generated using gene synthesis of a 984-bp fragment of *Spc25* RNAi resistant with 4bp *Drosophila* Kozak Sequence added in front of the start codon. The fragment was then cloned into XhoI site of vector PWG0773 pUAST-attB using sequence and ligation-independent cloning method. The pUAST-attB*-Spc25* RNAi resistant clone was used for microinjection into attP40 on 2L (25C7) (WellGenetics).

### Proximity ligation assay (PLA)

PLA is based on the following principle: Secondary antibodies conjugated with a PLA PLUS or PLA MINUS probe bind to anti-Flag and anti-Myc antibodies, respectively. During ligation, connector oligos hybridize to PLA probes and T4 ligases catalyze to form a circularized template. DNA polymerase amplifies the circularized template, which is bound by fluorescently labeled complementary oligos, allowing the interaction to be observed as PLA foci within the cells (Adopted from Duolink PLA, Merck). PLA was performed on S2 cells that were transfected with the following plasmids using Effectene Transfection Reagent (QIAGEN): control EGFP, control Flag, Flag-Arf1, EGFP-Ndc80, EGFP-Nuf2, Spc25-EGFP, Mis12-EGFP and Spc105-EGFP. The cells were washed thrice with cold PBS, fixed with 4% EM-grade formaldehyde in PBS for 15 min and blocked in 5% BSA in PBS-T (0.1% Triton-X100) for 45min. The cells were then incubated with primary antibodies at RT for 2 hrs before proceeding with Duolink PLA (Sigma-Aldrich) according to the manufacturer’s protocol. After incubation with primary antibodies, the cells were incubated with PLA probes at 37°C for 1 hr. They were then washed twice with Buffer A for 5 min, each at RT, followed by ligation of probes at 37°C for 30 min. Amplification was performed at 37°C for 100 min, followed by two washes with Buffer B, each for 10 min, at RT. The cells were washed once with 0.01x Buffer B before incubating with primary antibodies diluted in 3% BSA in PBS for 2 hrs at RT. Following this, the cells were washed twice with 0.1% PBS-T and incubated with secondary antibodies for 1.5 hrs at RT, before mounting them with in situ mounting media with DAPI (Duolink, Sigma-Aldrich).

### Brefeldin A (BFA)-treatment of *Drosophila* larval brains

Larval brains at 6h ALH were dissected in Shield and Sang M3 insect medium (Sigma-Aldrich) supplemented with 10% FBS. The brains were then incubated in either 10 μg/ml BFA (Sigma-Aldrich) dissolved in DMSO or the same volume of DMSO only as negative control in Shield and Sang M3 insect medium (Sigma-Aldrich) supplemented with 10% FBS. BFA treatment was carried out for 30 minutes. For Arf1 and GFP antibody detection after the treatments, the larval samples were fixed and immunostained as described above.

### Quantification and statistical analysis

*Drosophila* larval brains from various genotypes were placed dorsal side up on confocal slides. Confocal z-stacks were taken from the surface to the deep layers of the larval brains (20- 30 slides per z-stack with 2 or 3 µm intervals). For each genotype, all experiments were performed with a minimum of two biological replicates. In total a minimum of 6 brain lobes were imaged for z-stacks and Image J or Zen software’s was used for quantification.

Statistical analysis was performed using GraphPad Prism 8. Unpaired two-tailed t-tests were used for the comparison of two sample groups and one-way ANOVA or two-way ANOVA followed by Sidak’s multiple comparisons test were used for the comparison of more than two sample groups. All data are shown as mean ± SD. Statistically non-significant (ns), P > 0.05; * denotes P ≤ 0.05; ** denotes P ≤ 0.01; *** denotes P ≤ 0.001 and **** denotes P ≤ 0.0001.

### Data availability

All data generated or analyzed during this study are included in the manuscript. This study includes no data deposited in external repositories.

## Acknowledgements

We thank Christian F. Lehner, Gohta Goshima, C. Gonzalez, J. Raff, T. Lee, F. Matsuzaki, W. Chia, and the Bloomington *Drosophila* Stock Center, Vienna *Drosophila* Resource Center, Kyoto Stock Centre DGGR, and the Developmental Studies Hybridoma Bank for fly stocks and antibodies. We thank Drs Li Song and P.T. Ly for their involvement in the RNAi screening in the lab where *spc25* and *ndc80* RNAi phenotypes were identified. This work is supported by the Singapore National Research Foundation (NRF) Investigatorship (NRF-NRFI10-2024-0005) awarded to H. W. and the Ministry of Health, Singapore; National Medical Research Council, Singapore (MOH-001888) awarded to F.Y. and H.W.; NMRC Open Fund-Young Individual Research Grant (MOH-001514) awarded to M.G. Ministry of Education, Singapore under (MOE-T3-2020-01) and NRF (NRF-MSG-2023-0001) to YT.

## Author contributions

Conceptualization, MG and HW; Methodology, Data curation, and formal analysis, MG, HYA, GY, MDL, FY; Writing-original draft, MG and HW; Writing-review & editing, HW, MG, FY; funding acquisition, HW, FY, YT; Resources, HW, FY, YT; Supervision, HW.

## Supplementary Figure Legends

**Figure S1.**
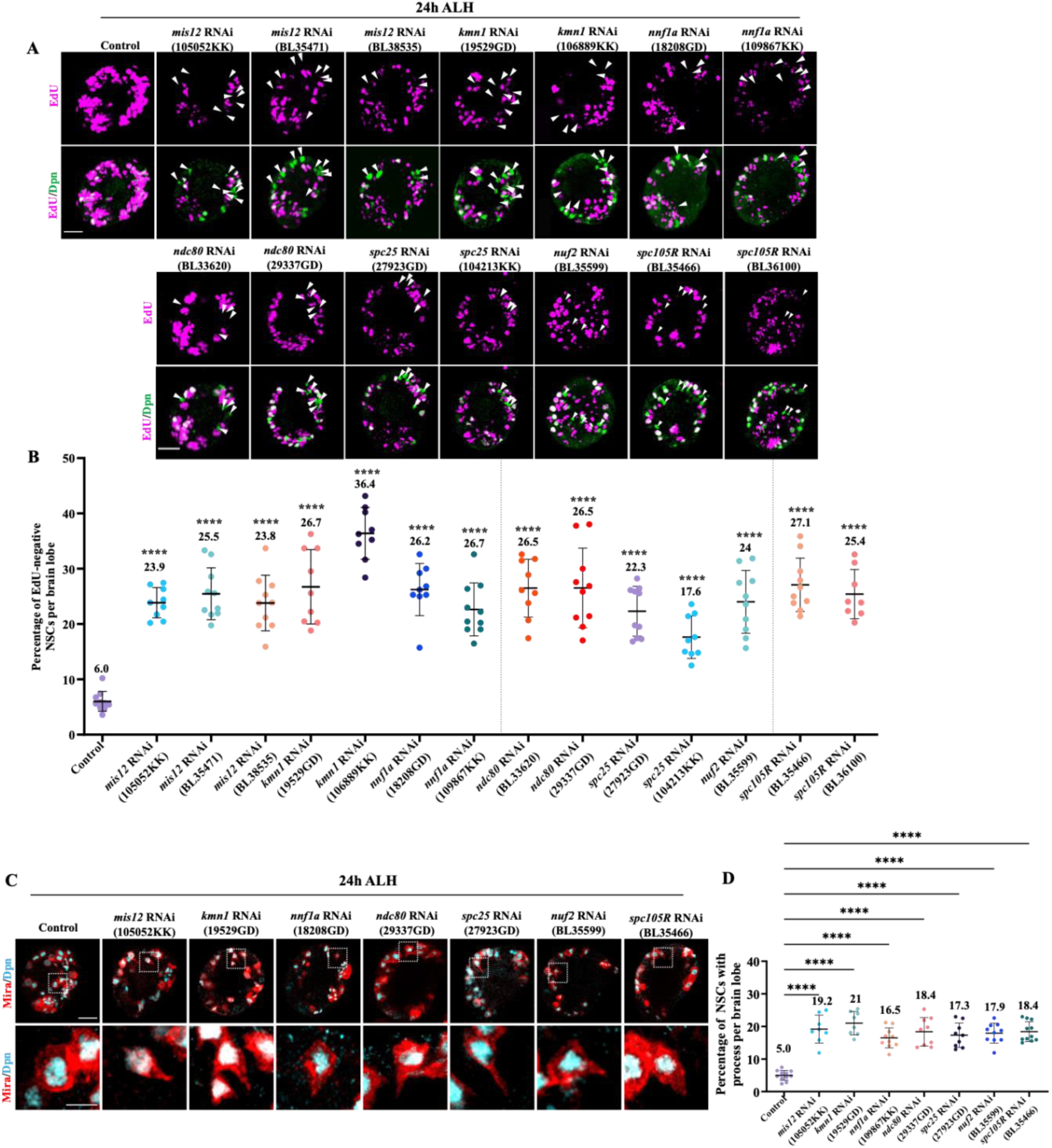
The kinetochore complex proteins are required for quiescent NSC reactivation. A) Larval brains at 24h ALH from control (*grh*-Gal4; UAS*-dicer2/*UAS*-β-Gal* RNAi), *mis12* RNAi (VDRC 105052KK), *mis12* RNAi (BDSC BL35471), *mis12* RNAi (BDSC BL38535), *kmn1* RNAi (VDRC 19529GD), *kmn1* RNAi (VDRC 106889KK), *nnf1a* RNAi (VDRC 18208GD), *nnf1a* RNAi (VDRC 109867KK), *ndc80* RNAi (BDSC BL33620), *ndc80* RNAi (VDRC 29337GD), *spc25* RNAi (VDRC 27923GD), *spc25* RNAi (VDRC 104213KK), *nuf2* RNAi (BDSC BL35599), *spc105R* RNAi (BDSC BL35466) and *spc105R* RNAi (BDSC BL36100) were analysed for EdU incorporation. NSCs were marked by Dpn. B) Quantification graph of EdU-negative NSCs per brain lobe for genotypes in (A). Control, 6%, n=10 BL; *mis12* RNAi (105052KK), 23.9%, n=9 BL; *mis12* RNAi (BL35471), 25.5%, n=10 BL; *mis12* RNAi (BL38535), 23.8%, n=10 BL; *kmn1* RNAi (19529GD), 26.7%, n=9 BL; *kmn1* RNAi (106889KK), 36.4%, n=9 BL; *nnf1a* RNAi (18208GD), 26.2%, n=9 BL; *nnf1a* RNAi (109867KK), 26.7%, n=10 BL; *ndc80* RNAi (BL33620), 26.5%, n=9 BL; *ndc80* RNAi (29337GD), 26.5%, n=10 BL; *spc25* RNAi (27923GD), 22.3%, n=10 BL; *spc25* RNAi (104213KK), 17.6%, n=9 BL; *nuf2* RNAi (BL35599), 24%, n=10 BL; *spc105R* RNAi (BL35466), 27.1%, n=10 BL; and *spc105R* RNAi (BL36100), 25.4%, n=8 BL. C) Larval brains at 24h ALH from control (*grh*-Gal4; UAS*-dicer2/*UAS*-β-Gal* RNAi), *mis12* RNAi (VDRC 105052KK), *kmn1* RNAi (VDRC 19529GD), *nnf1a* RNAi (VDRC 18208GD), *ndc80* RNAi (VDRC 29337GD), *spc25* RNAi (VDRC 27923GD), *nuf2* RNAi (BDSC BL35599) and *spc105R* RNAi (BDSC BL35466) were labelled with Dpn (nuclear marker) and Miranda (Mira) to mark qNSC protrusion. D) Quantification graph of NSCs retaining cellular protrusion per BL for genotypes in (A). Control, 5%, n=11 BL; *mis12* RNAi (105052KK), 19.2%, n=8 BL; *kmn1* RNAi (19529GD), 21%, n=8 BL; *nnf1a* RNAi (18208GD), 16.5%, n=10 BL; *ndc80* RNAi (29337GD), 18.4%, n=9 BL; *spc25* RNAi (27923GD), 17.3%, n=9 BL; *nuf2* RNAi (BL35599), 17.9%, n=10 BL and *spc105R* RNAi (BL35466), 18.4%, n=11 BL. Data information: (A-B) EdU incorporation was analyzed at 24h ALH by feeding larvae at 20h ALH with food supplemented with 0.2 mM EdU for 4h. White arrowheads point to NSCs without EdU incorporation. Data are presented as mean ± SD. Statistical significance was determined by one-way ANOVA with multiple comparisons. ns, non-significant, *p<0.05 and ****p<0.0001. Scale bars: 10μm.

**Figure S2.**
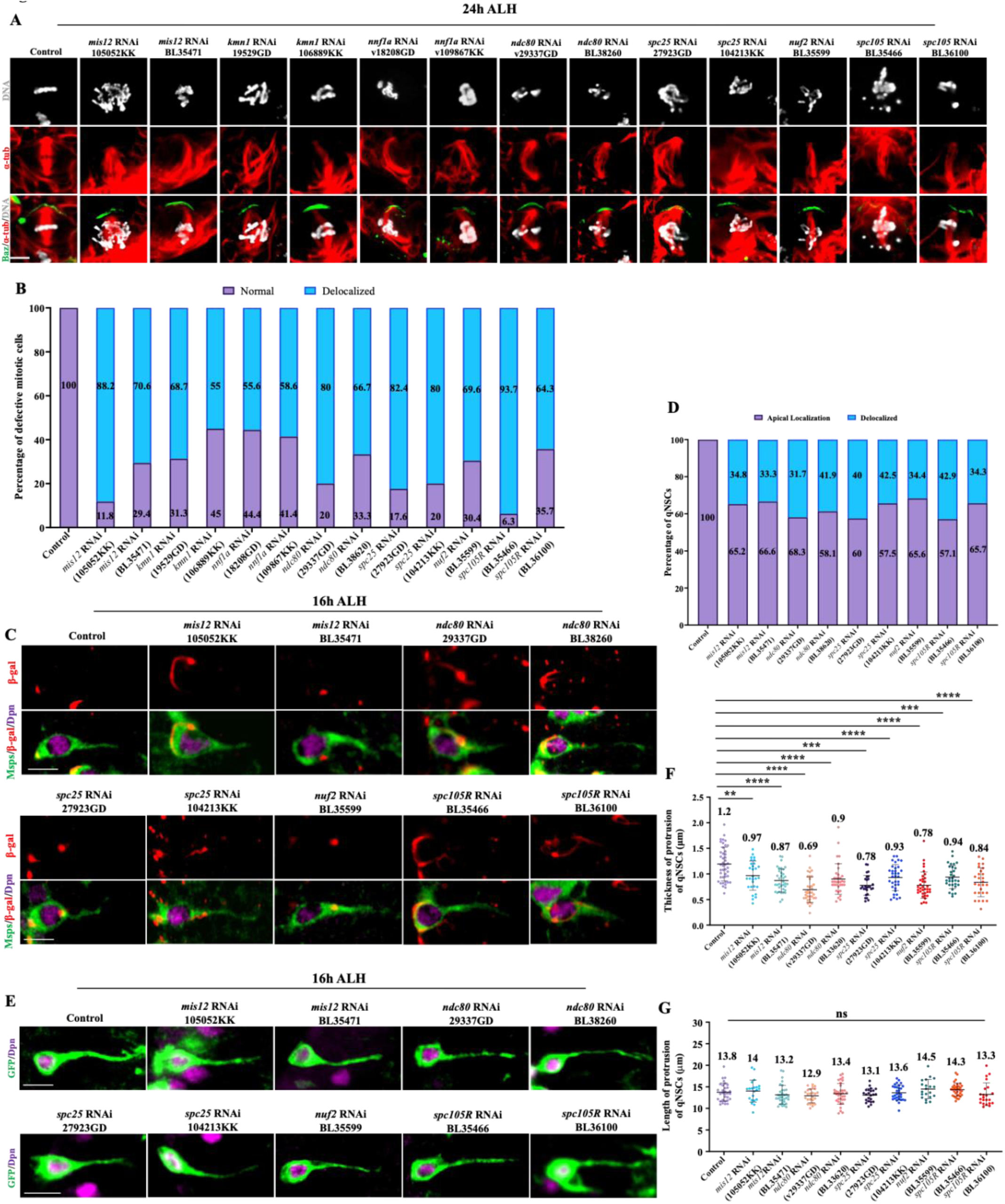
Kinetochore complex proteins are required microtubule orientation and morphological fidelity of qNSC cellular protrusion. A) Metaphase NSCs at 24h ALH from control (*grh*-Gal4; UAS*-dicer2/*UAS*-β-Gal* RNAi), *mis12* RNAi (VDRC 105052KK), *mis12* RNAi (BDSC BL35471), *kmn1* RNAi (VDRC 19529GD), *kmn1* RNAi (VDRC 106889KK), *nnf1a* RNAi (VDRC 18208GD), *nnf1a* RNAi (VDRC 109867KK), *ndc80* RNAi (VDRC 29337GD), *ndc80* RNAi (BDSC BL33620), *spc25* RNAi (VDRC 27923GD), *spc25* RNAi (VDRC 104213KK), *nuf2* RNAi (BDSC BL35599), *spc105R* RNAi (BDSC BL35466) and *spc105R* RNAi (BDSC BL36100) were stained for DNA, α-tubulin and Baz. B) Quantification graph of percentage of normal versus defective/de-localized DNA in metaphase NSCs for genotypes in (A). Control, Normal, 100%; Delocalized, 0% n=23 NSCs; *mis12* RNAi (105052KK), Normal, 11.8%; Delocalized, 88.2% n=17 NSCs; *mis12* RNAi (BL35471), Normal, 29.4%; Delocalized, 70.6% n=17 NSCs; *kmn1* RNAi (19529GD), Normal, 31.3%; Delocalized, 68.7% n=16 NSCs; *kmn1* RNAi (106889KK), Normal, 45%; Delocalized, 55% n=20 NSCs; *nnf1a* RNAi (18208GD), Normal, 44.4%; Delocalized, 55.6% n=27 NSCs; *nnf1a* RNAi (109867KK), Normal, 41.4%; Delocalized, 58.6% n=29 NSCs; *ndc80* RNAi (29337GD), Normal, 20%; Delocalized, 80% n=20 NSCs; *ndc80* RNAi (BL33620), Normal, 33.4%; Delocalized, 66.6% n=15 NSCs; *spc25* RNAi (27923GD), Normal, 17.6%; Delocalized, 82.4% n=17 NSCs; *spc25* RNAi (104213KK), Normal, 20%; Delocalized, 80% n=20 NSCs; *nuf2* RNAi (BL35599), Normal, 30.4%; Delocalized, 69.6% n=23 NSCs; *spc105R* RNAi (BL35466), Normal, 6.3%; Delocalized, 93.7% n=16 NSCs; and *spc105R* RNAi (BL36100), Normal, 35.7%; Delocalized, 64.3% n=14 NSCs. C) Larval brains at 16h ALH, including the control (UAS*-β-Gal* RNAi), *mis12* RNAi (105052KK), *mis12* RNAi (BL35471), *ndc80* RNAi (29337GD), *ndc80* RNAi (BL33620), *spc25* RNAi (27923GD), *spc25* RNAi (104213KK), *nuf2* RNAi (BL35599), *spc105R* RNAi (BL35466) and *spc105R* RNAi (BL36100) with Nod-β-Gal expressed under *insc*-Gal4, UAS-Dcr2 and *insc*-Gal4 were labelled with β-Gal, Dpn, and Msps. Quiescent NSCs at the central brain (CB) are shown. D) Quantification graph of Nod-β-Gal localization in qNSCs from genotypes in (C). De-localization of Nod-β-Gal from the apical region: control, 0%, n=24; *mis12* RNAi (105052KK), 34.8%, n=46; *mis12* RNAi (BL35471), 33.3%, n=27; *ndc80* RNAi (29337GD), 41.9%, n=43; *ndc80* RNAi (BL33620), 40%, n=30; *spc25* RNAi (27923GD), 42.5%, n=40; *spc25* RNAi (104213KK), 34.4%, n=32; *nuf2* RNAi (BL35599), 31.7%, n=41; *spc105R* RNAi (BL35466), 42.9%, n=35; and *spc105R* RNAi (BL36100), 34.3%, n=35. E) Larval brains at 16h ALH from control (UAS*-β-Gal* RNAi), *mis12* RNAi (105052KK), *mis12* RNAi (BL35471), *ndc80* RNAi (29337GD), *ndc80* RNAi (BL33620), *spc25* RNAi (27923GD), *spc25* RNAi (104213KK), *nuf2* RNAi (BL35599), *spc105R* RNAi (BL35466) and *spc105R* RNAi (BL36100) expressing *grh*>CD8-GFP were labeled with Dpn and GFP. F) Quantification graph for thickness of the primary protrusion of qNSCs from genotypes in (E). The thickness was measured at the middle point of the primary protrusion. Control, 1.2±0.33 μm, n=44 NSCs; *mis12* RNAi (105052KK), 0.97±0.27 μm, n=29 NSCs; *mis12* RNAi (BL35471), 0.87±0.25 μm, n=36 NSCs; *ndc80* RNAi (29337GD), 0.69±0.24 μm, n=35 NSCs; *ndc80* RNAi (BL33620), 0.9±0.3 μm, n=34 NSCs; *spc25* RNAi (27923GD), 0.78±0.19 μm, n=27 NSCs; *spc25* RNAi (104213KK), 0.93±0.25 μm, n=34 NSCs; *nuf2* RNAi (BL35599), 0.84±0.19 μm, n=27 NSCs; *spc105R* RNAi (BL35466), 0.78±0.19 μm, n=35 NSCs and *spc105R* RNAi (BL36100), 0.94±0.22μm, n=31 NSCs. G) Quantification graph for length of the primary protrusion of qNSCs for genotypes in (E). The length was measured from the PIS region of the primary protrusion to its neuropil contact site. Control, 13.8±2.1 μm, n=31 NSCs; *mis12* RNAi (105052KK), 14±2.5 μm, n=21 NSCs; *mis12* RNAi (BL35471), 13.2±2.2 μm, n=33 NSCs; *ndc80* RNAi (29337GD), 12.9±1.62 μm, n=25 NSCs; *ndc80* RNAi (BL33620), 13.4±2.4 μm, n=34 NSCs; *spc25* RNAi (27923GD), 13.1±1.5 μm, n=23 NSCs; *spc25* RNAi (104213KK), 13.6±1.75 μm, n=31 NSCs; *nuf2* RNAi (BL35599), 14.5±2.2 μm, n=20 NSCs; *spc105R* RNAi (BL35466), 14.3±1.5 μm, n=35 NSCs and *spc105R* RNAi (BL36100), 13.3±2.6 μm, n=23 NSCs. Data information: ns, non-significant, *p<0.05, **p<0.01, ***p<0.001 and ****p<0.0001 Data are presented as mean ± SD. Statistical significance was determined by one-way ANOVA with multiple comparisons. Scale bars: 5μm.

**Figure S3.**
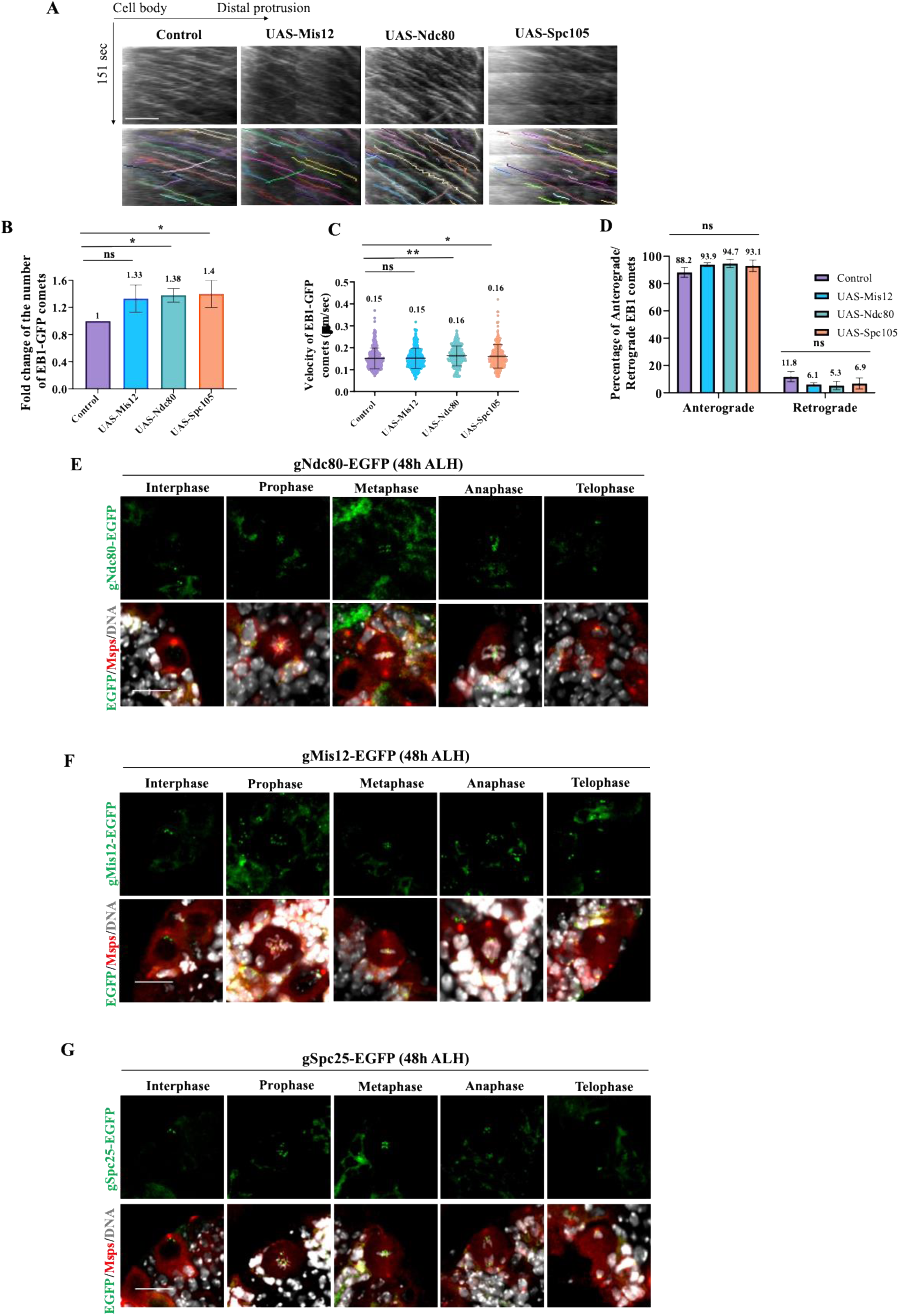
Kinetochore proteins are required for acentrosomal microtubule assembly in qNSCs. A) Kymographs of EB1-GFP comets movement in the primary protrusion of qNSCs expressing EB1-GFP under *grh*-Gal4 from the control, UAS*-*Mis12, UAS*-*Ndc80 and UAS*-*Spc105R at 6h ALH. The horizontal arrow indicates anterograde movement direction from cell body to the tip of the primary protrusion in qNSCs. B) Quantification graph of fold changes of number of EB1-GFP comets in the primary protrusion of qNSCs 6h ALH from various genotypes in (A). Control, 1, n=24 qNSCs, n=324 comets; UAS*-*Mis12, 1.33, n=17 qNSCs, n=310 comets; UAS*-*Ndc80, 1.38 n=14 qNSCs, n=263 comets and UAS*-*Spc105R, 1.4, n=15 qNSCs, n=281 comets. C) Quantification graph of velocity of EB1-GFP comets in the primary protrusion of qNSCs at 6h ALH from various genotypes in (A). Control, 0.15μm/sec, n=324 comets; UAS*-*Mis12, 0.15μm/sec, n=310 comets; UAS*-*Ndc80, 0.16μm/sec, n=263 comets and UAS*-*Spc105R, 0.16μm/sec, n=281 comets. D) Quantification graph of the percentage of anterograde- and retrograde-moving EB1-GFP comets in the primary protrusion of qNSCs from various genotypes in (A). Anterograde-moving comets: control, 88.2%, n=288 comets; UAS*-*Mis12, 93.9%, n=291 comets; UAS*-*Ndc80, 94.7%, n=249 comets and UAS*-*Spc105R, 93.1%, n=261 comets. Retrograde-moving comets: control, 11.8%, n=36 comets; UAS*-*Mis12, 6.1%, n=19 comets; UAS*-*Ndc80, 5.3%, n=14 comets and UAS*-*Spc105R, 6.9%, n=20 comets. E) Localization of genomic Ndc80-EGFP (gNdc80-EGFP) at 48h ALH during interphase, prophase, metaphase, anaphase and telophase. Cells were labelled for DNA and Msps. F) Localization of genomic Mis12-EGFP (gMis12-EGFP) at 48h ALH during interphase, prophase, metaphase, anaphase and telophase. Cells were labelled for DNA and Msps. G) Localization of genomic Spc25-EGFP (gSpc25-EGFP) at 48h ALH during interphase, prophase, metaphase, anaphase and telophase. Cells were labelled for DNA and Msps. Data information: ns, non-significant, *p<0.05, **p<0.01, ***p<0.001 and ****p<0.0001 Data are presented as mean ± SD. Statistical significance was determined by one-way ANOVA with multiple comparisons. Scale bars: 10μm for kymographs and 5μm for active NSCs.

**Figure S4.**
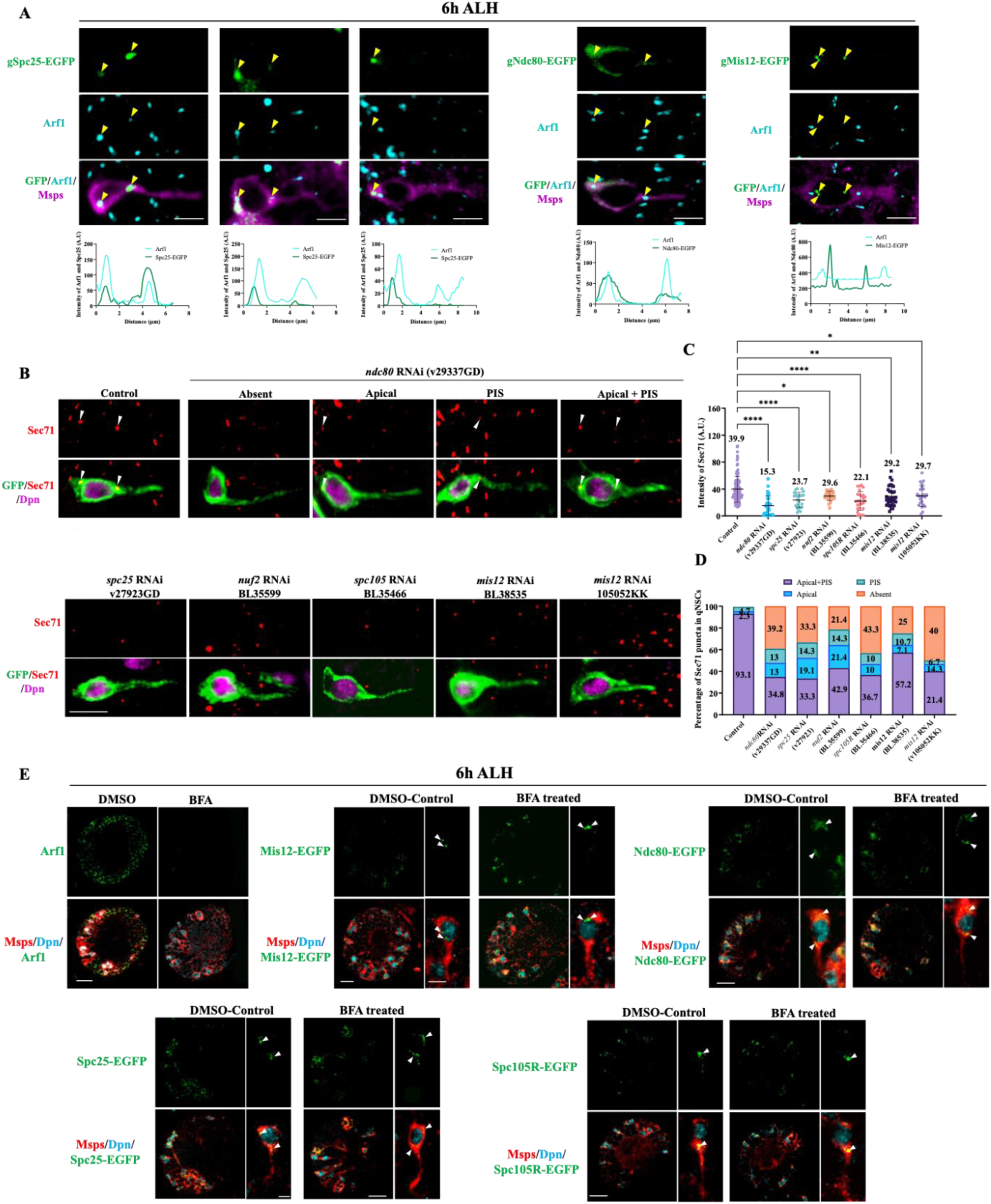
Kinetochore complex proteins are required for Arf1 and Sec71 localization at Golgi in qNSCs. A) Micrograph images of qNSCs at 6h ALH in gSpc25-EGFP, gNdc80-EGFP and gMis12-EGFP labelled with antibodies against GFP, Arf1, Msps and DNA. Line graphs show intensity curves of Arf1 (gray) with gSpc25-EGFP, gNdc80-EGFP and gMis12-EGFP (green) along the apical region to PIS region of the qNSC. Yellow arrowheads point to Kinetochore localization at, near or away from Arf1. B) qNSC with primary protrusions labelled by mCD8-GFP at 16h ALH from control (UAS*-β-Gal* RNAi), *ndc80* RNAi (29337GD), *spc25* RNAi (27923GD), *nuf2* RNAi (BL35599), *spc105R* RNAi (BL35466), *mis12* RNAi (BL38535) and *mis12* RNAi (105052KK) driven by the *gr*h-Gal4 driver, were labelled with antibodies against Sec71, Dpn, and GFP. C) Quantification graph of average intensity of Sec71 puncta for genotypes in (B). Control, 39.9 A.U., n=68; *ndc80* RNAi (29337GD), 15.3 A.U., n=32; *spc25* RNAi (27923GD), 23.7 A.U., n=24; *nuf2* RNAi (BL35599), 29.6 A.U., n=21; *spc105R* RNAi (BL35466), 22.1 A.U., n=27; *mis12* RNAi (BL38535), 29.2 A.U., n=32 and *mis12* RNAi (105052KK), 29.7 A.U., n=23. D) Quantification graph of the percentage of Sec71 puncta present per quiescent NSC for genotypes in (B). Control= Apical + PIS=93.1%, Apical=2.2%, PIS=4.7%, n=43; *ndc80* RNAi (29337GD), Apical + PIS=34.8%, Apical=13%, PIS=13%, Absent=39.2%, n=23; *spc25* RNAi (27923GD), Apical + PIS=33.3%, Apical=19.1%, PIS=14.3%, Absent=33.3%, n=21; *nuf2* RNAi (BL35599), Apical + PIS=42.9%, Apical=21.4%, PIS=14.3%, Absent=21.4%, n=14; *spc105R* RNAi (BL35466), Apical + PIS=36.7%, Apical=10%, PIS=10%, Absent=43.3%, n=30; *mis12* RNAi (BL38535), Apical + PIS=57.2%, Apical=7.1%, PIS=10.7%, Absent=25%, n=28 and *mis12* RNAi (105052KK), Apical + PIS=21.4%, Apical=14.3%, PIS=4.7%, Absent=40%, n=30. E) Larval brains at 6h ALH from Control, gMis12-EGFP, gNdc80-EGFP, gSpc25-EGFP and gSpc105R-EGFP were treated with DMSO and Brefeldin A (BFA) and stained for Arf1, GFP, Dpn and Msps to look at endogenous kinetochore localization in whole brain lobes and single qNSCs. Data information: Arrow heads indicate Sec71 puncta localization. Data are presented as mean ± SD. Statistical significance was determined by one-way ANOVA with multiple comparisons. *p<0.05, **p<0.01, ***p<0.001 and ****p<0.0001. Scale bars: 10μm for whole BLs and 5μm for single qNSCs.

**Figure S5.**
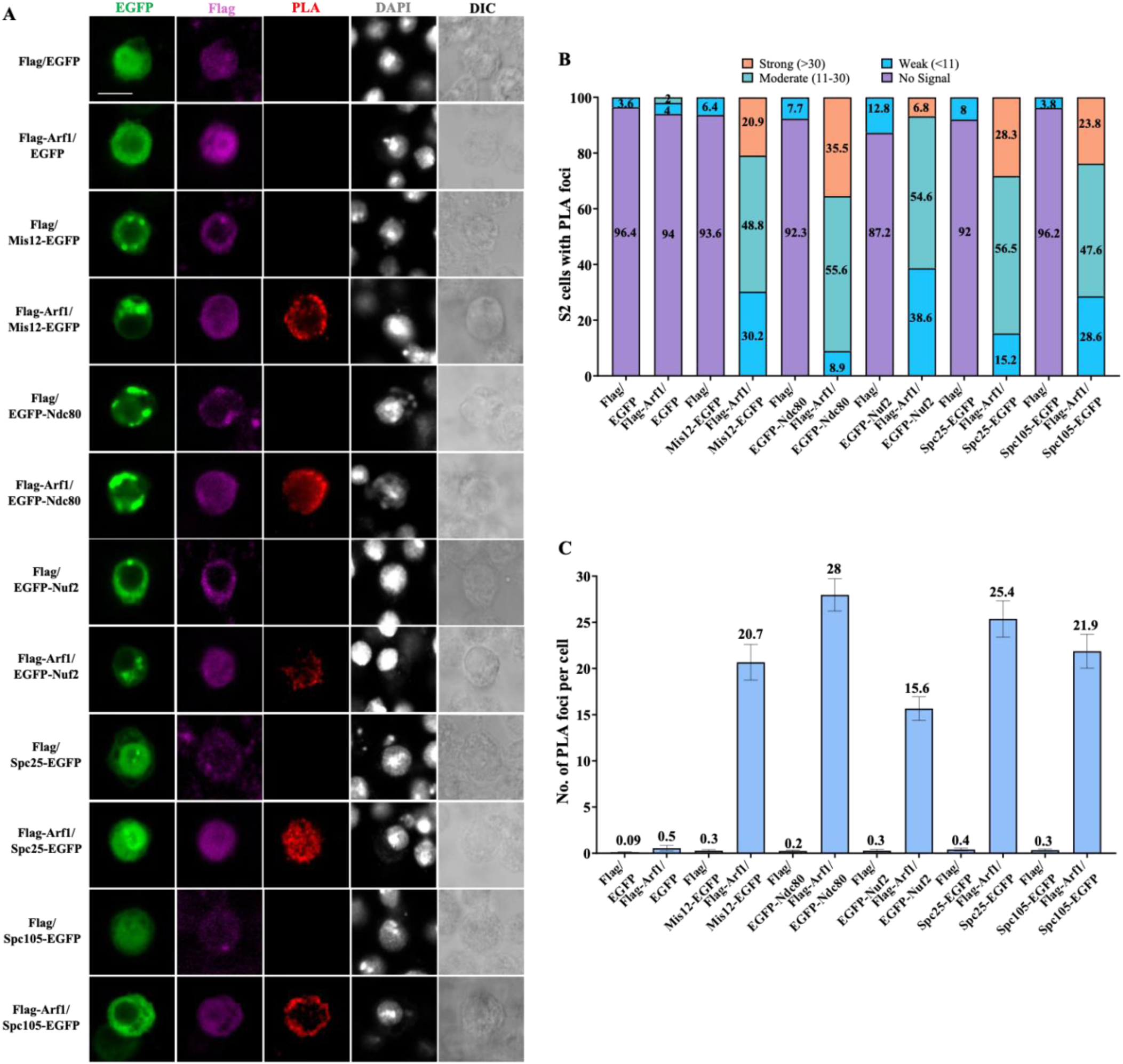
Kinetochore complex proteins physically associate with the Golgi protein Arf1. A) *In situ* PLA between Arf1^WT^ and Mis12-EGFP, EGFP-Ndc80, EGFP-Nuf2, Spc25-EGFP or Spc105-EGFP in S2 cells. S2 cells transfected with two of the indicated plasmids (Flag, GFP, Flag-Arf1^WT^, Mis12-EGFP/ EGFP-Ndc80/ EGFP-Nuf2/ Spc25-EGFP or Spc105-EGFP). S2 cells were then stained for Flag, GFP, and DNA and detected for PLA signal. Cell outlines were determined by differential interference contrast (DIC) images. B) Quantification graphs showing the percentage of S2 cells with no PLA signal, weak (≤10 foci), moderate (11–30 foci), and strong (>30 foci) PLA signals for (A). Flag and EGFP, no signal=96.4%, weak signal=3.6%, n=112; Flag-Arf1^WT^ and EGFP, no signal=94%, weak=4%, moderate=2%, n=100; Flag and Mis12-EGFP, no signal=93.6%, weak=6.4%, n=94; Flag-Arf1^WT^ and Mis12-EGFP, no signal=0%, weak=30.2%, moderate=48.8%, strong=20.9%, n=86; Flag and EGFP-Ndc80, no signal=92.3%, weak=7.7%, n=104; Flag-Arf1^WT^ and EGFP-Ndc80, no signal=0%, weak=8.9%, moderate=55.6%, strong=35.5%, n=90; Flag and EGFP-Nuf2, no signal=87.2%, weak=12.8%, n=94; Flag-Arf1^WT^ and EGFP-Nuf2, no signal=0%, weak=38.6%, moderate=54.6%, strong=6.8%, n=88; Flag and Spc25-EGFP, no signal=92%, weak=8%, n=100; Flag-Arf1^WT^ and Spc25-EGFP, no signal=0%, weak=15.2%, moderate=56.5%, strong=28.3%, n=92 and Flag and Spc105-EGFP, no signal=96.2%, weak=3.8%, n=106; Flag-Arf1^WT^ and Spc105-EGFP, no signal=0%, weak=28.6%, moderate=47.6%, strong=23.8%, n=84. C) Quantification graph of the average number of PLA foci per cell in (B). Flag and EGFP, 0.09, n=112; Flag-Arf1^WT^ and EGFP, 0.5, n=100; Flag and Mis12-EGFP, 0.3, n=94; Flag-Arf1^WT^ and Mis12-EGFP, 20.7, n=86; Flag and EGFP-Ndc80, 0.2, n=104; Flag-Arf1^WT^ and EGFP-Ndc80, 28, n=90; Flag and EGFP-Nuf2, 0.3, n=94; Flag-Arf1^WT^ and EGFP-Nuf2, 15.6, n=88; Flag and Spc25-EGFP, 0.4, n=100; Flag-Arf1^WT^ and Spc25-EGFP, 25.4, n=92 and Flag and Spc105-EGFP, 0.3, n=106; Flag-Arf1^WT^ and Spc105-EGFP, 21.9, n=84. Data information: Data are presented as mean ± SD. Scale bars: 5μm.

**Figure S6.**
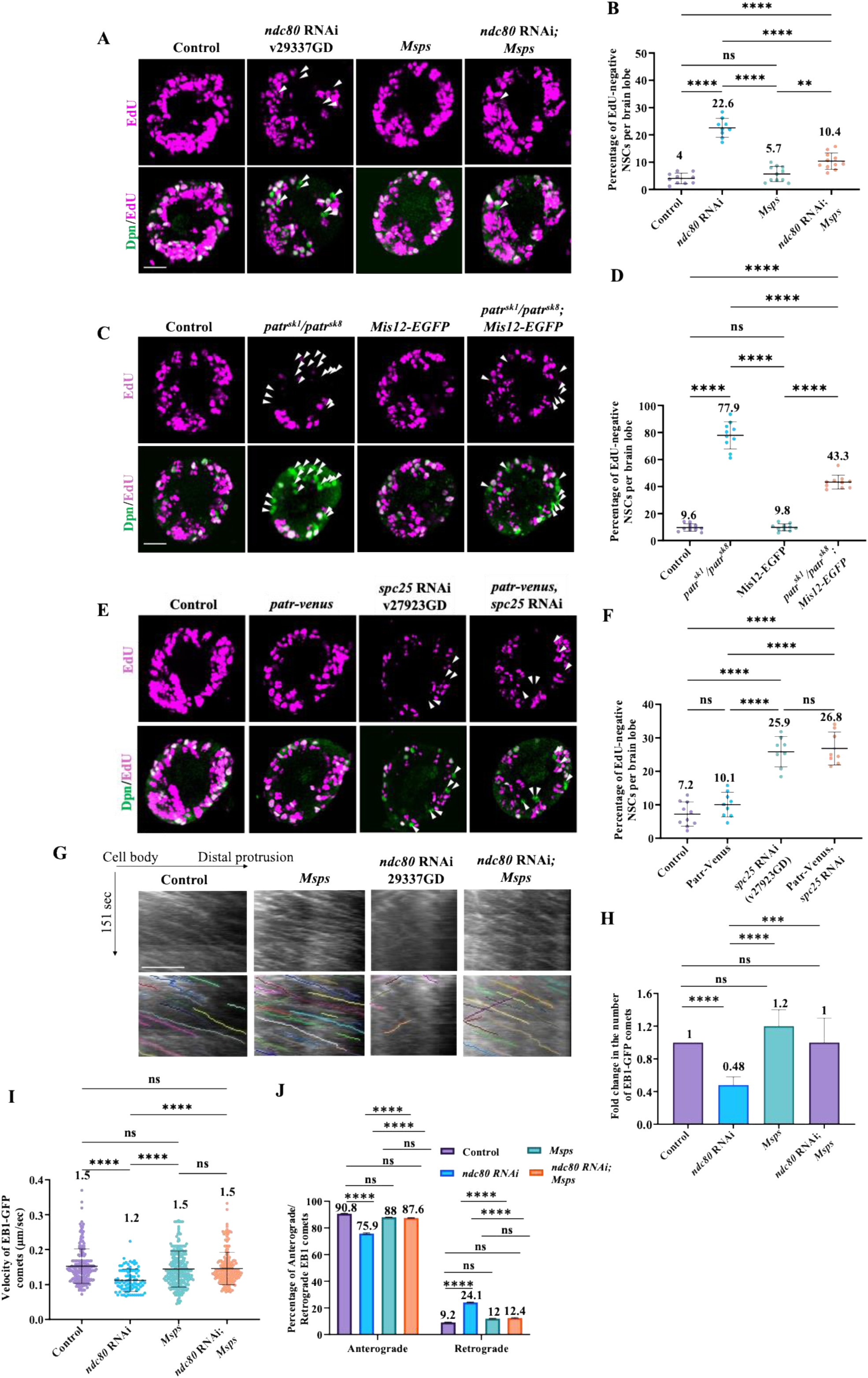
Kinetochore proteins function upstream of Msps and downstream of Patronin to promote NSC reactivation. A) Larval brains at 24h ALH from control (UAS*-β-Gal* RNAi), *ndc80* RNAi (29337GD), UAS-Msps and *ndc80* RNAi (29337GD); UAS-Msps under *grh*-Gal4 were analysed for EdU incorporation. NSCs were marked by Dpn. B) Quantification graph of EdU-negative NSCs per brain lobe for genotypes in (A). Control, 4%, n=10 BL; *ndc80* RNAi (29337GD), 22.6%, n=9 BL; UAS-Msps, 5.7%, n=11 BL and *ndc80* RNAi (29337GD); UAS-Msps, 10.4%, n=11 BL. C) Larval brains at 24h ALH from control (*UAS-β-Gal* RNAi), *patr^sk1^/patr^sk8^*, *UAS-*Mis12-EGFP and *patr^sk1^/patr^sk8^*; UAS*-*Mis12-EGFP under *grh*-Gal4 were analysed for EdU incorporation. NSCs were marked by Dpn. D) Quantification graph of EdU-negative NSCs per brain lobe for genotypes in (C). Control, 9.6%, n=10 BL; *patr^sk1^/patr^sk8^*, 77.9%, n=10 BL; UAS*-*Mis12-EGFP, 9.8%, n=10 BL and *patr^sk1^/patr^sk8^*; UAS*-*Mis12-EGFP, 43.3%, n=10 BL. E) Larval brains at 24h ALH from control (UAS*-β-Gal* RNAi), UAS*-*Patr-Venus, *spc25* RNAi (27923GD) and *spc25* RNAi (27923GD); UAS*-*Patr-Venus was analysed for EdU incorporation. NSCs were marked by Dpn. F) Quantification graph of EdU-negative NSCs per brain lobe for genotypes in (E). Control, 7.2%, n=10 BL; UAS*-*Patr-Venus, 10.1%, n=9 BL; *spc25* RNAi (27923GD), 25.9%, n=8 BL and *spc25* RNAi (27923GD); UAS*-*Patr-Venus, 26.8%, n=8 BL. G) Kymographs of EB1-GFP comets movement in the primary protrusion of qNSCs expressing EB1-GFP under *grh*-Gal4 from the control, *ndc80* RNAi (29337GD), UAS-Msps and *ndc80* RNAi (29337GD); UAS-Msps at 6h ALH. The horizontal arrow indicates anterograde movement direction from cell body to the tip of the primary protrusion in qNSCs. H) Quantification graph of fold changes of number of EB1-GFP comets in the primary protrusion of qNSCs 6h ALH from various genotypes in (G). Control, 1, n=16 qNSCs, n=207 comets; *ndc80* RNAi (29337GD), 0.48, n=15 qNSCs, n=87 comets; UAS-Msps, 1.2 n=15 qNSCs, n=233 comets and *ndc80* RNAi (29337GD); UAS-Msps, 1, n=15 qNSCs, n=201 comets. I) Quantification graph of velocity of EB1-GFP comets in the primary protrusion of qNSCs at 6h ALH from various genotypes in (G). Control, 0.15μm/sec, n=207 comets; *ndc80* RNAi (29337GD), 0.12μm/sec, n=87 comets; UAS-Msps, 0.15μm/sec, n=233 comets and *ndc80* RNAi (29337GD); UAS-Msps, 0.15μm/sec, n=201 comets. J) Quantification graph of the percentage of anterograde- and retrograde-moving EB1-GFP comets in the primary protrusion of qNSCs from various genotypes in (G). Anterograde-moving comets: control, 90.8%, n=188 comets; *ndc80* RNAi (29337GD), 75.9%, n=66 comets; UAS-Msps, 88%, n=176 comets *ndc80* RNAi (29337GD); UAS-Msps, 87.6%, n=205 comets. Retrograde-moving comets: control, 9.2%, n=19 comets; *ndc80* RNAi (29337GD), 24.1%, n=21 comets; UAS-Msps, 12%, n=28 comets and *ndc80* RNAi (29337GD); UAS-Msps, 12.4%, n=25 comets. Data information: (A-F) EdU incorporation was analyzed at 24h ALH by feeding larvae at 20h ALH with food supplemented with 0.2 mM EdU for 4h. White arrowheads point to NSCs without EdU incorporation. Data are presented as mean ± SD. Statistical significance was determined by one-way ANOVA with multiple comparisons. ns, non-significant, *p<0.05, **p<0.01, ***p<0.001 and ****p<0.0001. Scale bars: 10μm.

**Figure S7.**
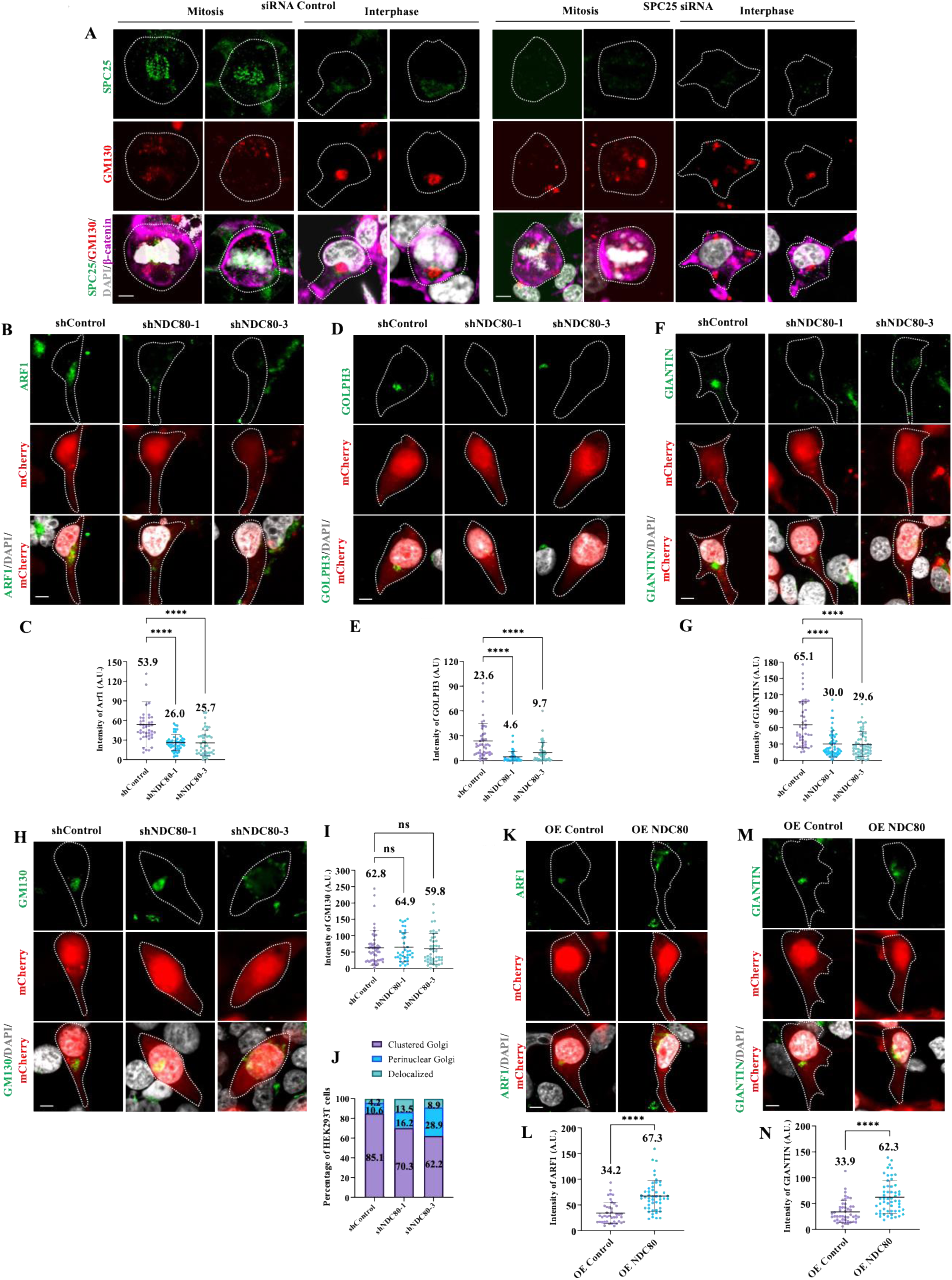
Kinetochore protein NDC80 is required for Golgi localization in HEK293T cells. A) Immunostaining micrographs showing antibody specificity of human SPC25 in mitotic and interphase HEK293T cells. SPC25 was knocked down in HEK293T cells by siRNA. siRNA against SPC25 showed a strong loss of SPC25 puncta at mitotic stages and reduced or no cytoplasmic localization in interphase cells. Cells are labelled by SPC25 (green), GM130 (red), DAPI (gray) and β-catenin (magenta, to mark cell outline). siControl, N=64; siSPC25, N=79. B) Immunostaining micrographs showing ARF1 localization in shControl, shNDC80-1 and shNDC80-3 in HEK293T cells 72h after transfection. Transfected cells are labelled by endogenous mCherry. DNA is stained by DAPI. C) Quantification graph of average intensity of ARF1 for genotypes in (B). shControl, 53.9 A.U., n=42; shNDC80-1, 26 A.U., n=50; shNDC80-3, 25.7 A.U., n=44. D) Immunostaining micrographs showing GOLPH3 localization in shControl, shNDC80-1 and shNDC80-3 in HEK293T cells 72h after transfection. Transfected cells are labelled by endogenous mCherry. DNA is stained by DAPI. E) Quantification graph of average intensity of GOLPH3 for genotypes in (D). shControl, 23.6 A.U., n=45; shNDC80-1, 4.6 A.U., n=39; shNDC80-3, 9.7 A.U., n=44. F) Immunostaining micrographs showing GIANTIN localization in shControl, shNDC80-1 and shNDC80-3 in HEK293T cells 72h after transfection. Transfected cells are labelled by endogenous mCherry. DNA is stained by DAPI. G) Quantification graph of average intensity of GIANTIN for genotypes in (F). shControl, 65.1 A.U., n=45; shNDC80-1, 30 A.U., n=51; shNDC80-3, 29.6 A.U., n=57. H) Immunostaining micrographs showing GM130 localization in shControl, shNDC80-1 and shNDC80-3 in HEK293T cells 72h after transfection. Transfected cells are labelled by endogenous mCherry. DNA is stained by DAPI. I) Quantification graph of average intensity of GM130 for genotypes in (H). shControl, 62.8 A.U., n=47; shNDC80-1, 64.9 A.U., n=37; shNDC80-3, 57.8 A.U., n=45. J) Quantification graph of condensed, dispersed or delocalized GM130 for genotypes in (H). shControl, Clustered Golgi= 85.1%, Perinuclear Golgi=10.6%, De-localized Golgi=4.2%, n=47; shNDC80-1, Clustered Golgi= 70.3%, Perinuclear Golgi=16.2%, De-localized Golgi=13.5%, n=37; Clustered Golgi= 62.2%, Perinuclear Golgi=28.9%, De-localized Golgi=8.9%, n=45. K) Immunostaining micrographs showing ARF1 localization in OE Control and OE NDC80 in HEK293T cells 72h after transfection. Transfected cells are labelled by endogenous mCherry. DNA is stained by DAPI. L) Quantification graph of average intensity of ARF1 for genotypes in (K). OE Control, 34.2 A.U., n=41; OE NDC80, 67.3 A.U., n=48. M) Immunostaining micrographs showing GIANTIN localization in OE Control and OE NDC80 in HEK293T cells 72h after transfection. Transfected cells are labelled by endogenous mCherry. DNA is stained by DAPI. N) Quantification graph of average intensity of GIANTIN for genotypes in (M). OE Control, 33.9 A.U., n=47; OE NDC80, 62.4 A.U., n=55. Data information: ns, non-significant, *p<0.05, **p<0.01, ***p<0.001 and ****p<0.0001 Data are presented as mean ± SD. For (A-I), Statistical significance was determined by one-way ANOVA with multiple comparisons. For (J-M), Statistical significance was determined by unpaired student’s T-test. Scale bars: 10μm.

**Figure S8.**
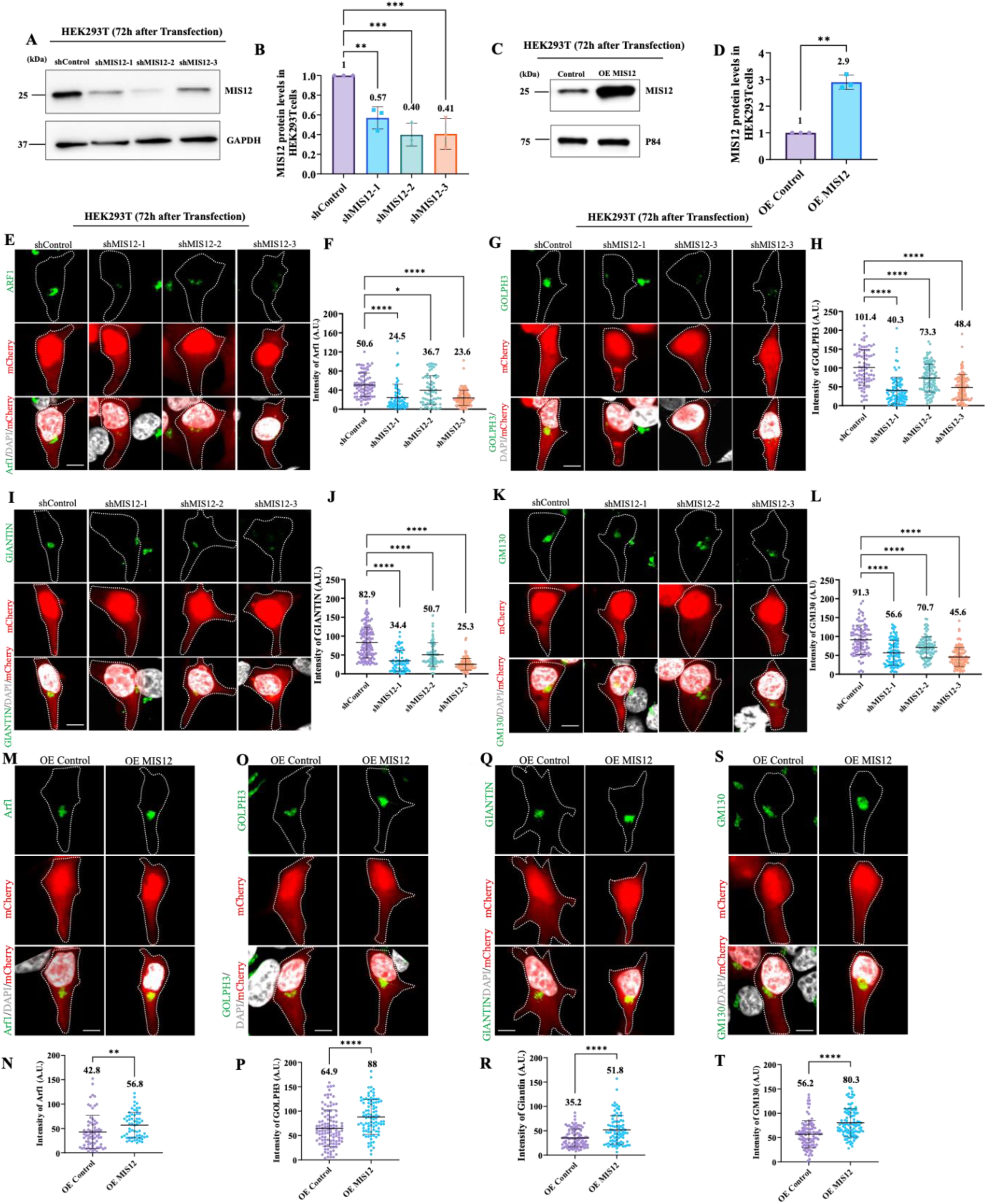
Kinetochore protein MIS12 is required for Golgi localization in HEK293T cells. A) WB analysis of HEK293T cell protein extracts from control (shControl-mCherry) and three different MIS12 KDs (shMIS12-1, shMIS12-2 and shMIS12-3) with lentivirus infection in 72h culture. Blots were probed with anti-MIS12 antibody and anti-GAPDH antibody. Immunoblot showing the efficiency of MIS12 KD. B) Quantification graph representing fold change in knockdown efficiency of MIS12 normalized to the internal control GAPDH and shControl. shControl, 1, n=3; shMIS12-1, 0.57, n=3; shMIS12-2, 0.40, n=3; shMIS12-3, 0.41, n=3. C) WB analysis of HEK293T cell protein extracts from control (Over expression (OE) Control-mCherry) and OE MIS12 with lentivirus infection in 72h culture. Blots were probed with anti-MIS12 antibody and anti-GAPDH antibody. Immunoblot showing the efficiency of MIS12over expression. D) Quantification graph representing fold change in over expression efficiency of MIS12 normalized to the internal control GAPDH and OE Control. OE Control, 1, n=3; OE MIS12, 2.9, n=3. E) Immunostaining micrographs showing ARF1 localization in shControl, shMIS12-1, SHMIS12-2 and shMIS12-3 in HEK293T cells 72h after transfection. Transfected cells are labelled by endogenous mCherry. DNA is stained by DAPI. F) Quantification graph of average intensity of ARF1 for genotypes in (E). shControl, 50.6 A.U., n=81; shMIS12-1, 24.5 A.U., n=64; shMIS12-2, 36.7 A.U., n=66 AND shMIS12-3, 23.6 A.U., n=102. G) Immunostaining micrographs showing GOLPH3 localization in shControl, shMIS12-1, SHMIS12-2 and shMIS12-3 in HEK293T cells 72h after transfection. Transfected cells are labelled by endogenous mCherry. DNA is stained by DAPI. H) Quantification graph of average intensity of GOLPH3 for genotypes in (G). shControl, 101.4 A.U., n=84; shMIS12-1, 40.3 A.U., n=94; shMIS12-2, 73.3 A.U., n=107 and shMIS12- 3, 48.4 A.U., n=106. I) Immunostaining micrographs showing GIANTIN localization in shControl, shMIS12-1, SHMIS12-2 and shMIS12-3 in HEK293T cells 72h after transfection. Transfected cells are labelled by endogenous mCherry. DNA is stained by DAPI. J) Quantification graph of average intensity of GIANTIN for genotypes in (I). shControl, 82.9 A.U., n=139; shMIS12-1, 34.4 A.U., n=64; shMIS12-2, 50.7 A.U., n=70 and shMIS12-3, 25.3 A.U., n=107. K) Immunostaining micrographs showing GM130 localization in shControl, shMIS12-1, SHMIS12-2 and shMIS12-3 in HEK293T cells 72h after transfection. Transfected cells are labelled by endogenous mCherry. DNA is stained by DAPI. L) Quantification graph of average intensity of GM130 for genotypes in (K). shControl, 91.3 A.U., n=98; shMIS12-1, 56.6 A.U., n=98; shMIS12-2, 70.7 A.U., n=91 and shMIS12-3, 45.6 A.U., n=103. M) Immunostaining micrographs showing ARF1 localization in OE Control and OE MIS12 in HEK293T cells 72h after transfection. Transfected cells are labelled by endogenous mCherry. DNA is stained by DAPI. N) Quantification graph of average intensity of ARF1 for genotypes in (M). OE Control, 42.8 A.U., n=71; OE MIS12, 56.8 A.U., n=60. O) Immunostaining micrographs showing GOLPH3 localization in OE Control and OE MIS12 in HEK293T cells 72h after transfection. Transfected cells are labelled by endogenous mCherry. DNA is stained by DAPI. P) Quantification graph of average intensity of GOLPH3 for genotypes in (O). OE Control, 64.9 A.U., n=108; OE MIS12, 88 A.U., n=91. Q) Immunostaining micrographs showing GIANTIN localization in OE Control and OE MIS12 in HEK293T cells 72h after transfection. Transfected cells are labelled by endogenous mCherry. DNA is stained by DAPI. R) Quantification graph of average intensity of GIANTIN for genotypes in (Q). OE Control, 35.2 A.U., n=94; OE MIS12, 51.8 A.U., n=83. S) Immunostaining micrographs showing GM130 localization in OE Control and OE MIS12 in HEK293T cells 72h after transfection. Transfected cells are labelled by endogenous mCherry. DNA is stained by DAPI. T) Quantification graph of average intensity of GM130 for genotypes in (S). OE Control, 56.2 A.U., n=98; OE MIS12, 80.3 A.U., n=102. Data information: ns, non-significant, *p<0.05, **p<0.01, ***p<0.001 and ****p<0.0001 Data are presented as mean ± SD. For (B, E-L), Statistical significance was determined by one-way ANOVA with multiple comparisons. For (C, M-T), Statistical significance was determined by unpaired student’s T-test. Scale bars: 10μm.

**Figure S9.**
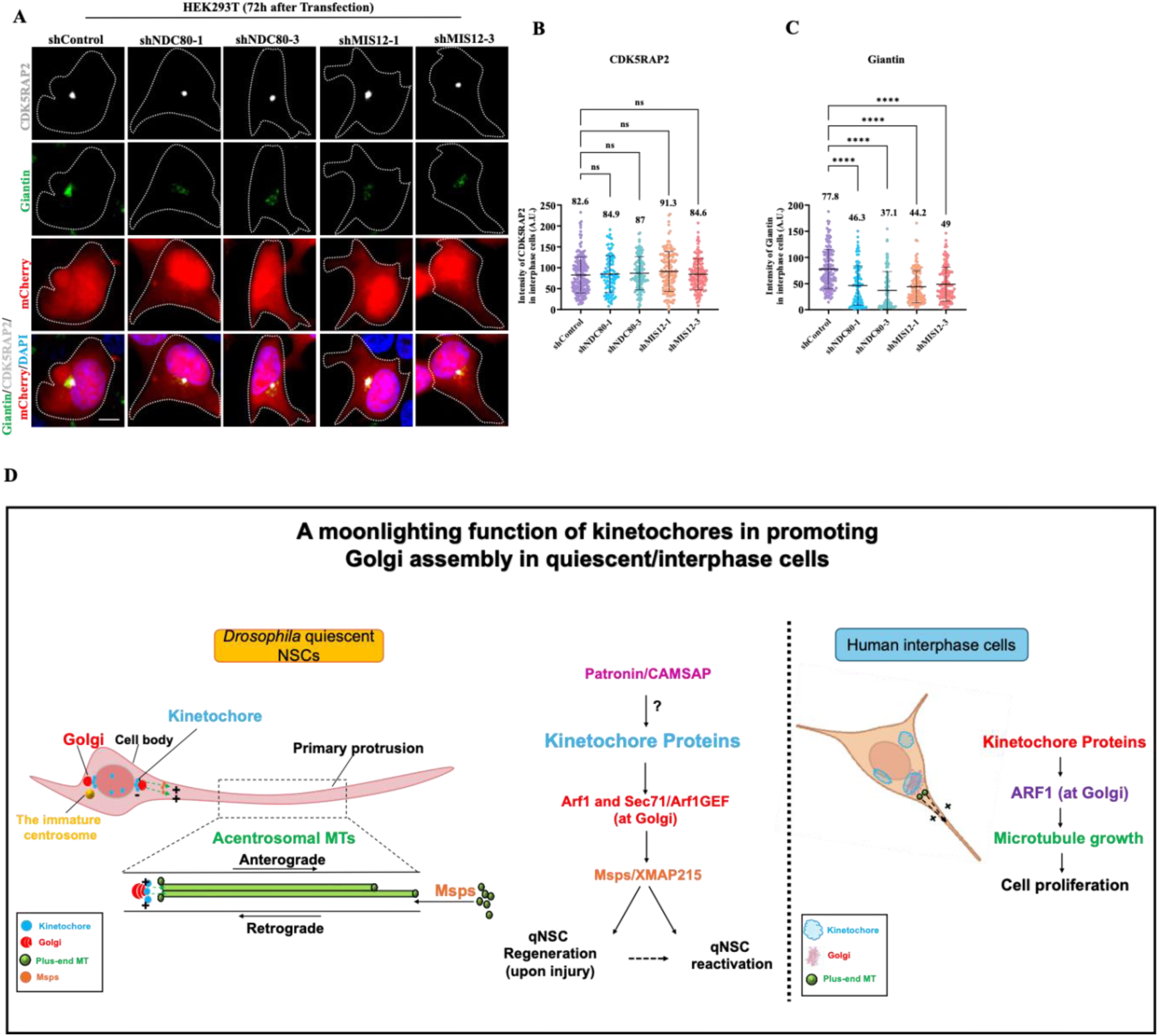
Kinetochore proteins are not required for centrosomal localization in interphase HEK293T cells. A) Immunostaining micrographs showing CDK5RAP2 and GIANTIN localization in shControl, shNDC80-1, shNDC80-3, shMIS12-1 and shMIS12-3 in interphase HEK293T cells 72h after transfection. Transfected cells are labelled by endogenous mCherry. DNA is stained by DAPI. B) Quantification graph of average intensity of CDK5RAP2 for genotypes in (A). shControl, 82.6 A.U., n=190; shNDC80-1, 84.9 A.U., n=101; shNDC80-3, 87 A.U., n=116; shMIS12-1, 91.3 A.U., n=135; and shMIS12-3, 84.6 A.U., n=129. C) Quantification graph of average intensity of GIANTIN for genotypes in (A). shControl, 77.8 A.U., n=152; shNDC80-1, 46.3 A.U., n=99; shNDC80-3, 37.1 A.U., n=98; shMIS12-1, 44.2 A.U., n=122; and shMIS12-3, 49 A.U., n=123. D) Working Model: In *Drosophila* quiescent NSCs, the primary protrusion is supported by a highly polarized microtubule network that is predominantly acentrosomal and oriented with plus ends facing outward toward the neuropil. Kinetochore proteins regulate Golgi organization and positioning by controlling the localization of Golgi-associated factors, including the small GTPase Arf1, in quiescent NSCs. Kinetochore proteins physically associate with Arf1 to regulate acentrosomal microtubule growth and quiescent NSC reactivation. Kinetochore proteins act upstream of Arf1 and the microtubule polymerase Msps/XMAP215 and downstream of the microtubule minus-end binding protein Patronin/CAMSAP to regulate acentrosomal microtubule assembly during quiescent NSC regeneration and reactivation. In human interphase cells, kinetochore proteins localize near or at the Golgi and are required for proper Golgi organization and microtubule growth dynamics. Data information: ns, non-significant, *p<0.05, **p<0.01, ***p<0.001 and ****p<0.0001 Data are presented as mean ± SD. For (B-C) Statistical significance was determined by one-way ANOVA with multiple comparisons. For (E-H), Statistical significance was determined by unpaired student’s T-test. Scale bars: 10μm.

## Supplementary Movies

**Movie S1.** Time-lapse imaging of EB1-GFP comets in the primary protrusion of a quiescent NSC in the larval brain expressing *grh*-Gal4; UAS-EB1-GFP with UAS-*β-Gal* RNAi, *mis12* RNAi (105052KK), *kmn1* RNAi (19529GD), *nnf1a* RNAi (18202GD), *ndc80* RNAi (BL33620), *spc25* RNAi (104213KK), *nuf2* RNAi (BL35599) and *spc105R* RNAi (BL35466) at 6h ALH. Time scale: minute: second. Scale bar: 10 µm.

**Movie S2.** Time-lapse imaging of EB1-GFP comets in the primary protrusion of a quiescent NSC in the larval brain expressing *grh*-Gal4; UAS-EB1-GFP with UAS-*β-Gal* RNAi, UAS-Mis12, UAS-Ndc80 and UAS-Spc105R at 6h ALH. Time scale: minute: second. Scale bar: 10 µm.

**Movie S3.** Time-lapse imaging of regeneration of the primary protrusion of a quiescent NSC expressing *grh*-Gal4; UAS-mCD8-GFP with UAS-β-Gal RNAi, *mis12* RNAi (105052KK), *nuf2* RNAi (BL35599), *spc25* RNAi (27923GD) and *spc105R* RNAi (BL35466) in *ex vivo* larval brain at 12h ALH after laser ablation. Time scale: minute: second. Scale bar: 10 µm.

**Movie S4.** Time-lapse imaging of EB1-GFP comets in the primary protrusion of a quiescent NSC in the larval brain expressing *grh*-Gal4; UAS-EB1-GFP with UAS-*β-Gal* RNAi, UAS-Arf1, *spc25* RNAi (27923GD) and Arf1; *spc25* RNAi (27923GD) at 6h ALH. Time scale: minute: second. Scale bar: 10 µm.

**Movie S5.** Time-lapse imaging of EB1-GFP comets in the primary protrusion of a quiescent NSC in the larval brain expressing *grh*-Gal4; UAS-EB1-GFP with UAS-*β-Gal* RNAi, UAS-Msps, *ndc80* RNAi (29337GD) and *ndc80* RNAi (29337GD); UAS*-* Msps at 6h ALH. Time scale: minute: second. Scale bar: 10 µm.

**Movie S6.** Time-lapse imaging of regeneration of the primary protrusion of a quiescent NSC expressing *grh*-Gal4; UAS-mCD8-GFP with UAS-*β-Gal* RNAi, UAS-Arf1, *spc25* RNAi (27923GD) and Arf1; *spc25* RNAi (27923GD) in *ex vivo* larval brain at 12h ALH after laser ablation. Time scale: minute: second. Scale bar: 10 µm.

**Movie S7** Time-lapse imaging tracking the growing ends of microtubules by using the plus-end microtubule binding protein EB3 tagged with Tdtomato in mouse NPCs in shControl and shArf1. Time scale: minute: second. Scale bar: 10 µm.

**Movie S8.** Time-lapse imaging tracking the growing ends of microtubules by using the plus-end microtubule binding protein EB3 tagged with GFP in human NPCs in shControl and shNDC80. Time scale: minute: second. Scale bar: 10 µm.

**Movie S9** Time-lapse imaging tracking the growing ends of microtubules by using the plus-end microtubule binding protein EB3 tagged with GFP in human NPCs in OE Control and OE NDC80. Time scale: minute: second. Scale bar: 10 µm.

